# SWI/SNF antagonizes SIR heterochromatin to promote transcription of genes expressed during mitotic exit in *Saccharomyces cerevisiae*

**DOI:** 10.1101/2020.03.24.006205

**Authors:** Mayuri Rege, Jessica L. Feldman, Nicholas L. Adkins, Craig L. Peterson

## Abstract

Heterochromatin is a repressive, specialized chromatin structure that is central to eukaryotic transcriptional regulation and genome stability. In the budding yeast, *Saccharomyces cerevisiae*, heterochromatin formation requires Sir2p, Sir3p, and Sir4p, and these Sir proteins create specialized chromatin structures at telomeres and silent mating type loci. Previously, we reported that the SWI/SNF chromatin remodeling enzyme can evict Sir3 from chromatin fibers *in vitro*, though whether this activity contributes to the role of SWI/SNF as a transcriptional activator at euchromatic loci is unknown. Here, we characterize genetic interactions between the *SIR* genes (*SIR2*, *SIR3*, and *SIR4*) and genes encoding subunits of the chromatin remodelers SWI/SNF and INO80C, as well genes encoding the histone deacetylases Hst3 and Hst4. We find that loss of *SIR* genes partially rescues the growth defects of *swi2*, *ino80*, and *hst3/hst4* mutants during replication stress conditions. Interestingly, partial suppression of *swi2*, *ino80*, and *hst3 hst4* mutant phenotypes is due to the pseudo-diploid state of *sir* mutants, but a significant portion is due to more direct functional interactions. Consistent with this view, transcriptional profiling of strains lacking Swi2 or Sir3 identifies a set of genes whose expression in the M/G1 phase of the cell cycle requires SWI/SNF to antagonize the repressive impact of Sir3.

## INTRODUCTION

Eukaryotic genomes are packaged with positively charged histone proteins to form chromatin. Chromatin can be divided into two functional categories: transcriptionally active euchromatin and transcriptionally silent heterochromatin. In budding yeast, heterochromatic structures are formed at each telomere and the two silent mating type loci (*HMR* and *HML*). The assembly of heterochromatin domains requires the binding of non-histone proteins to the chromatin fiber, and in yeast these are Sir2, Sir3 and Sir4 (Rusche *et al*. 2003; Rusché *et al*. 2002). Sir3 is believed to be the structural component of yeast heterochromatin, whereas Sir2 is a histone deacetylase that functions with Sir4 to target Sir proteins to the proper genomic locations. Deletion of *SIR* genes leads to expression of both a- and α-specific genes in haploids, producing a pseudo-diploid state (Haber 1998). Previous studies have found that this pseudo-diploid state alters the DNA damage response, alleviating the genotoxic stress phenotypes of several *rad* mutants (Schild 1995; Valencia-Burton *et al*. 2006).

In addition to heterochromatic regions, Sir3 has also been detected by chromatin immunoprecipitation studies at euchromatic locations, although the functional implications are not understood (Radman-Livaja *et al*. 2011). Immunofluorescence studies of Sir3 have also revealed that Sir3 forms discrete nuclear puncta for most of the cell cycle, except for a diffuse nuclear staining pattern during mitotic stages (Laroche *et al*. 2000). Overexpression of Sir3 can lead to the expansion of heterochromatin domains and gene-silencing defects within euchromatin (Taddei *et al*. 2009; Holmes *et al*. 1997), indicating that aberrant binding of Sir3 to euchromatic sites can be detrimental.

ATP-dependent chromatin remodeling enzymes are a major contributor to the dynamic nature of chromatin. They modify chromatin structure by mobilizing or disrupting nucleosomes in an ATP-dependent reaction (Clapier and Cairns 2009). The SWI/SNF chromatin remodeling enzyme is a founding member of this group of enzymes (Smith and Peterson 2005), and subunits of SWI/SNF were first identified in yeast genetic studies as global activators of transcription (Peterson *et al*. 1992; Laurent *et al*. 1991). For instance, inactivation of the Swi2 ATPase subunit leads to defects in the transcription of many inducible yeast genes, as well the function of many transcriptional activators. Strains harboring mutations in genes encoding SWI/SNF subunits have severe growth defects, are sensitive to DNA damaging or replication stress agents, and show a defect in mitotic exit (Krebs *et al*. 2000). Similar to other remodeling enzymes, SWI/SNF can use the energy of ATP hydrolysis to mobilize nucleosomes in cis, or evict nucleosomal H2A/H2B dimers as well as entire histone octamers from DNA. Recently, we also found that SWI/SNF has the novel ability to catalyze the displacement of the Sir3 protein from nucleosomal substrates *in vitro*. This activity is not shared with other remodeling enzymes, and it requires a direct interaction between the Swi2 subunit and Sir3. This activity appears to be important for cells to contend with replication stress (Manning and Peterson 2014).

In this work, we report genetic interactions between the gene encoding the Swi2 subunit of SWI/SNF and genes encoding the Sir2, Sir3, and Sir4 heterochromatin components. Inactivation of Sir3 alleviated the slow growth phenotype of a *swi2Δ* strain, and partially restored resistance to the replication stress agent, hydroxyurea (HU). Deletion of *SIR2* or *SIR3* also partially suppressed the replication stress phenotypes caused by loss of the INO80C chromatin remodeling complex as well as the loss of the H3 lysine 56-specific histone deacetylases Hst3 and Hst4. Interestingly, in some cases partial suppression of genotoxic stress phenotypes were observed in pseudo-diploid cells, suggesting indirect as well as direct impacts of Sir3 loss. To identify potential transcriptional targets for the SWI/SNF-Sir3 antagonism, we characterized the transcriptional profile of *swi2Δ*, *sir3Δ*, and *swi2Δ sir3Δ* strains. A parallel analysis was also performed in *SIR3* and *sir3Δ* strains where Swi2 was conditionally depleted from the nucleus by the anchor away method (Haruki *et al*. 2008). This latter method circumvented transcriptional defects due to the severe growth phenotype of the *swi2Δ* strain, and together identified a common set of genes where SWI/SNF promotes transcription by antagonizing Sir3.

## MATERIALS AND METHODS

### Yeast growth media and genetic methods

Yeast were cultured using standard procedures (Rege *et al*. 2015). For tetrad analysis, at least 30 tetrads were dissected for segregation analysis and growth rates noted.

## List of strains

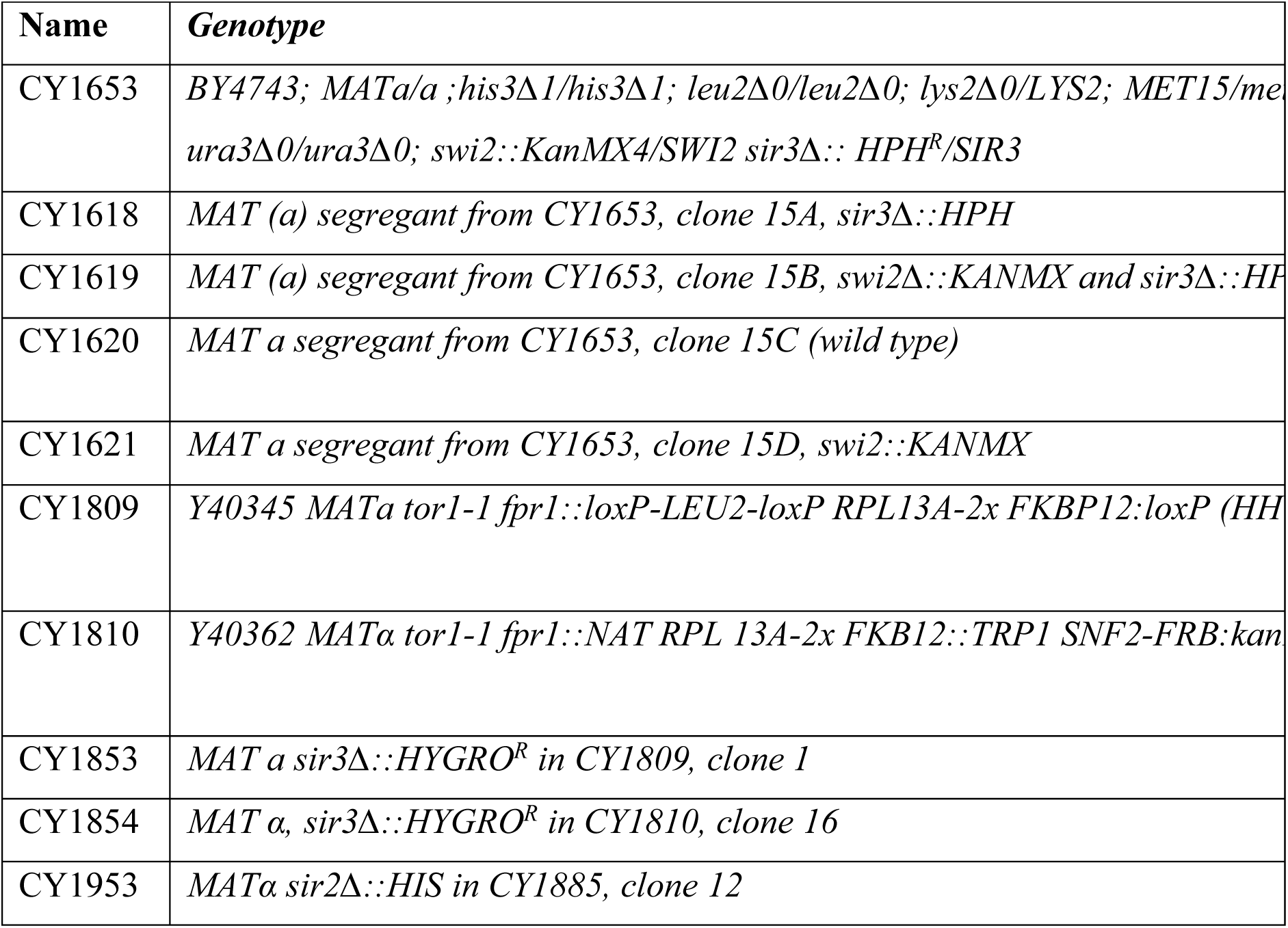
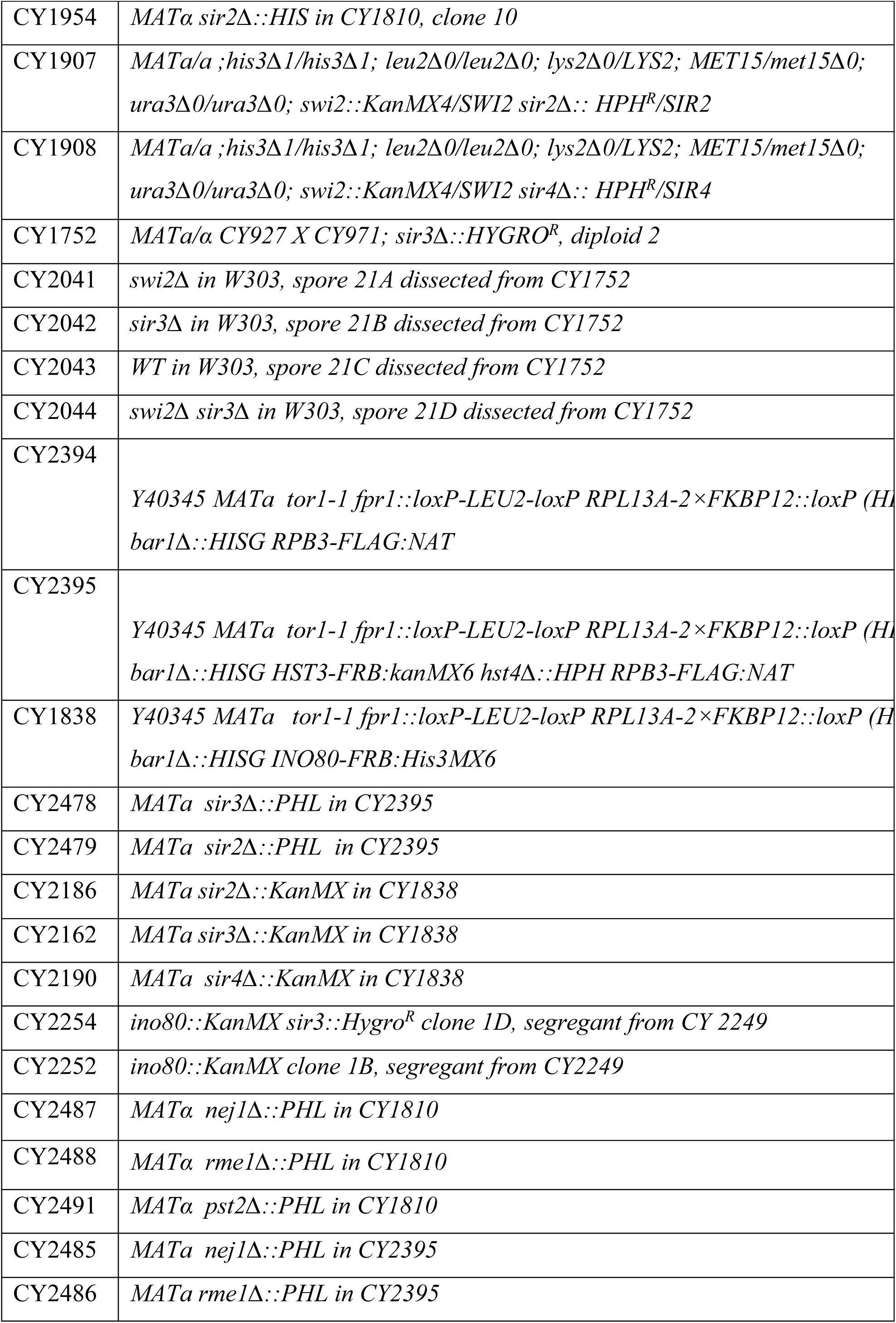
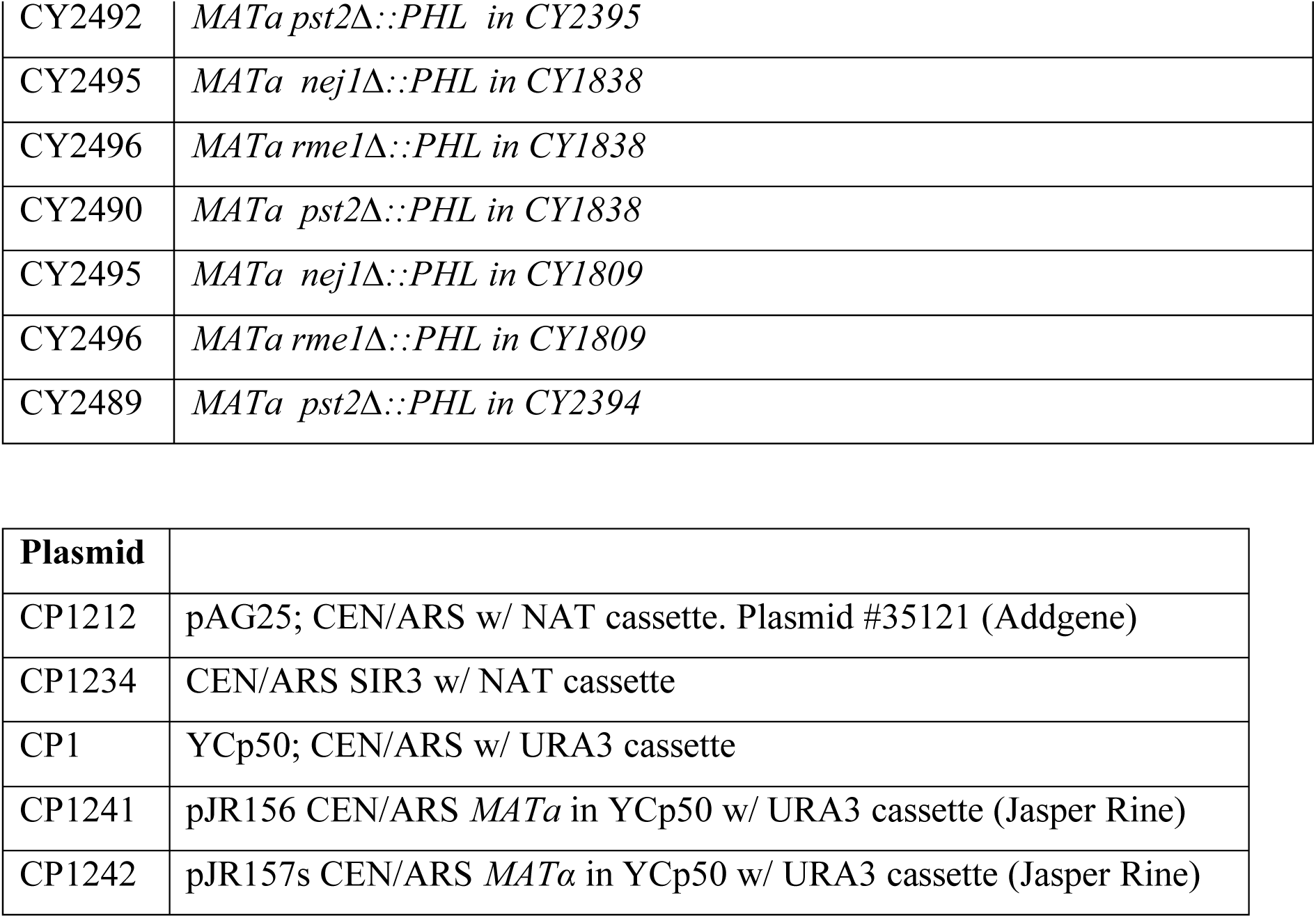

### Chromatin Immunoprecipitation (ChIP)

Yeast strains were grown in rich media with 2% glucose at 30°C and either DMSO or Rapamycin (8µg/ml final concentration) was added for 60 minutes before fixation with 1.2% formaldehyde. Cells were quenched with 2.5M glycine, centrifuged, rinsed with cold water and stored at −80°C until chromatin preparation. Chromatin preparation, immunoprecipitation and DNA extraction were performed as described in (Bennett *et al*. 2013). The anti-Sir3 antibody (1 µL for 100µL chromatin) was used to immunoprecipitate native Sir3. The anti-H3 antibody, ab1791 from Abcam (1 µL for 100µL chromatin) was used to immunoprecipitate histone H3. The *SIR3* gene was C-terminally tagged with a FLAG tag and an anti-FLAG antibody used for immunoprecipitation.

### Microarray sample preparation and analysis

Yeast strains were grown in rich media with 2% glucose at 30°C in 50 ml cultures, collected at OD = 0.8 for RNA preparation and RNA was extracted using the hot phenol method as described previously (Rege *et al*. 2015). Samples prepared as described in Welch *et al*. 2007 were hybridized to Affymetrix Yeast 2.0 arrays from four replicates of *swi2*Δ and *swi2*Δ *sir3*Δ strains and analyzed by limma analysis in R (Bioconductor package). Yeast strains were grown in rich media with 2% glucose at 30°C to OD = 0.6. and either DMSO or Rapamycin (8µg/ml final concentration) was added for 60 minutes and pelleted for RNA preparation (Rege *et al*. 2015). One replicate each of the *SWI2-FRB*, *SWI2-FRB sir3Δ* and *sir3Δ* arrays and corresponding WT arrays was used. Total RNA was hybridized on Affymetrix Yeast 2.0 arrays and analyzed using a log2 fold change cut-off. The raw data files have been deposited on the GEO database (# in process).

### qRT-PCR

Samples for total RNA were prepared and qRT-PCR was performed as described previously in Manning and Peterson 2014.

### Data Availability Statement

All strains made in this study are available upon request from the Peterson lab. The lists of Group 1_KO genes and Group 1_AA genes are given in Table S1 and Table S2, respectively. The list of Group 1 genes common in the KO and AA datasets are given in Table S3. The lists of Group 2_KO genes and Group 2_AA genes are given in Table S4 and Table S5, respectively. RMA normalized data obtained using GeneSpring Affymetrix Software for all the conditions and replicates are provided in Tables S5 and S6. Raw microarray. CEL files have been deposited in NCBI’s GEO database with the accession number (in process).

## RESULTS

### The slow growth phenotype of *swi2*Δ is partially rescued by *sir3*Δ

An isogenic set of wildtype, *sir3Δ, swi2Δ*, and *swi2Δ sir3Δ* strains was created by tetrad dissection from a swi2Δ/*SWI2 sir3Δ/SIR3* heterozygous diploid. Deletion of *SIR3* partially suppresses the growth defect of *swi2*Δ on rich media (Figure 1A), suggesting that these loci genetically interact. Importantly, this suppression segregates with markers for the double mutant after tetrad analysis, eliminating the possibility that a nonspecific, background suppressor causes the growth suppression in *swi2Δ sir3Δ* strains (Figure 1A). We also find that the growth defects of *swi2Δ* are suppressed by *sir3Δ* in a different strain background (w303; Figure 1B). In addition to slow growth on glucose media, *swi2*Δ mutants are unable to metabolize alternative carbon sources like raffinose, galactose, glycerol, or ethanol (Abrams *et al*. 1986; Carlson *et al*. 1981). Inactivation of Sir3 did not facilitate growth of a *swi2Δ* on raffinose, but limited suppression was observed for growth on media containing galactose, ethanol, or glycerol (Figure 1C). *SWI2* is also required for resistance to replication stress, induced by hydroxyurea (HU), and a *swi2*Δ shows a delayed growth rate in this condition (Sharma *et al*. 2003). Interestingly, deletion of *SIR3* partially relieves the HU sensitive phenotype of a *swi2*Δ (Figure 1C). Thus, a subset of *swi2Δ* phenotypes are alleviated by deletion of *SIR3*.

**Figure 1:**
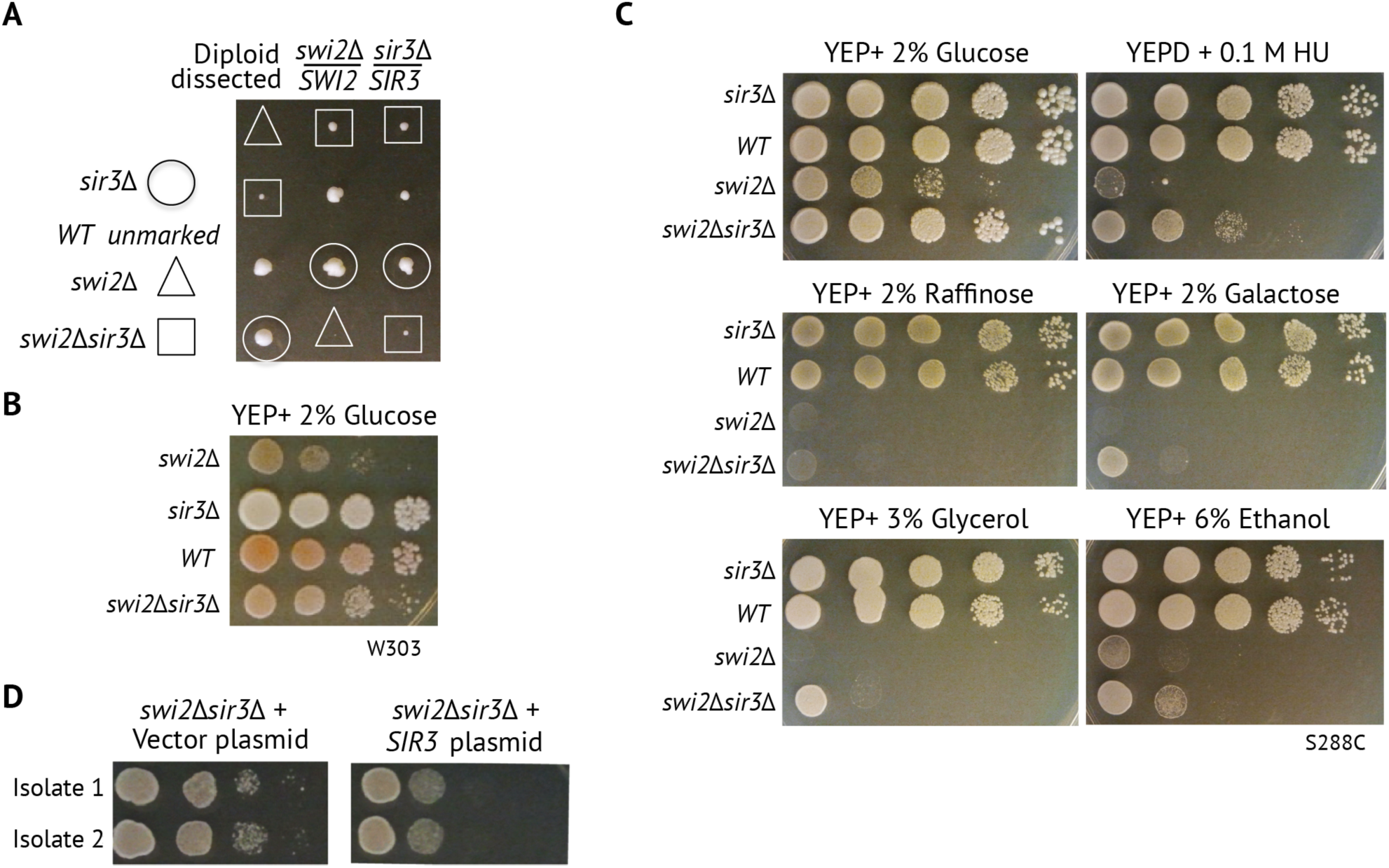
*swi2* growth defects are partially rescued by deletion of *SIR3*. A) Tetrad dissection plates of the *swi2Δ/SWI2 sir3Δ/SIR3* heterozygous diploid on YEPD plates with the corresponding genotypes marked with symbols listed on the left. A single dissected spore yields an isogenic colony, imaged after 10 days. Relative size of each colony is representative of the growth rate. B) Spot assay on null mutants dissected from the W303 background. Equal cell numbers were spotted in consecutive ten-fold dilutions on agar plates with 2% glucose as the carbon source and imaged after 3 days. C) Spot assay was performed as described in B) with different carbon sources. Raffinose and galactose plates also contain 2% antimycin to prevent respiratory growth. D) *swi2Δ sir3Δ* mutants transformed with a plasmid containing either the vector backbone (left) or with a construct expressing Sir3 from its endogenous promoter (right). Spot assays were performed on individual isolates as described in B).

To completely eliminate the possibility that a background mutation other than the *sir3Δ* segregated with, and caused the growth suppression seen in the double mutant, we transformed the *swi2Δ sir3Δ* with a plasmid containing *SIR3* expressed from its endogenous promoter. As expected, complementation with a vector plasmid had no impact on growth, while the *SIR3* plasmid slowed the growth of the *swi2Δ sir3Δ* strain (Figure 1D). Given that *sir3Δ* suppresses the severe growth defects of *swi2Δ* in multiple strain backgrounds, and that this suppression can be reversed when *swi2Δ sir3Δ* is complemented by a *SIR3* plasmid, these data suggest that SWI/SNF antagonizes Sir3 *in vivo*.

### Absence of *SIR2* does not suppress *swi2*Δ growth defects

Given that *SIR3* shows negative genetic interactions with *SWI2*, we asked whether genes that encode other Sir proteins, Sir2 and Sir4, also showed similar genetic interactions. Sir2 is a histone deacetylase (HDAC) that promotes Sir3 binding to nucleosomes by removing the acetyl group on histone H4 lysine 16. Sir4 forms a complex with Sir2, and it is believed to play a key role in targeting Sir proteins to telomeres and HM loci (Rusché *et al*. 2002; Thurtle and Rine 2014). Unlike deletion of *SIR3*, inactivation of Sir2 did not alleviate the slow growth of the *swi2Δ* (Figure S1A, B). In contrast, inactivation of Sir4 suppresses the growth defect of a *swi2Δ* mutant (Figure S1C, D). Thus, the studies support genetic interactions between *SWI2*, *SIR3*, and *SIR4*, but not *SIR2*.

### Comparison of *swi2Δ* alleles with conditional depletion of Swi2

As *swi2Δ* null mutants are extremely slow growing, we wanted to establish an alternative approach to interrogate the genetic interactions between *SIR* genes and *SWI2*. To this end, the anchor away system was used to conditionally deplete Swi2 from the nucleus (Haruki *et al*. 2008). The parent strain harbors a <u>FK</u>506 <u>b</u>inding <u>p</u>rotein (FKBP12) tag fused to the C-terminus of an anchor protein, RPL13A. RPL13A is a ribosomal protein that is present in high copy numbers in the cell and transits from the nucleus to the cytoplasm during ribosome assembly, as shown in Figure 2A. In this parent strain, we tagged the endogenous *SWI2* locus at the C-terminus with the FKBP12-rapamycin-binding (FRB) domain. Rapamycin induces formation of a ternary complex between the FKBP12 and FRB domains, and thus, rapidly depletes *SWI2*-FRB from the nucleus (Figure 2A).

**Figure 2:**
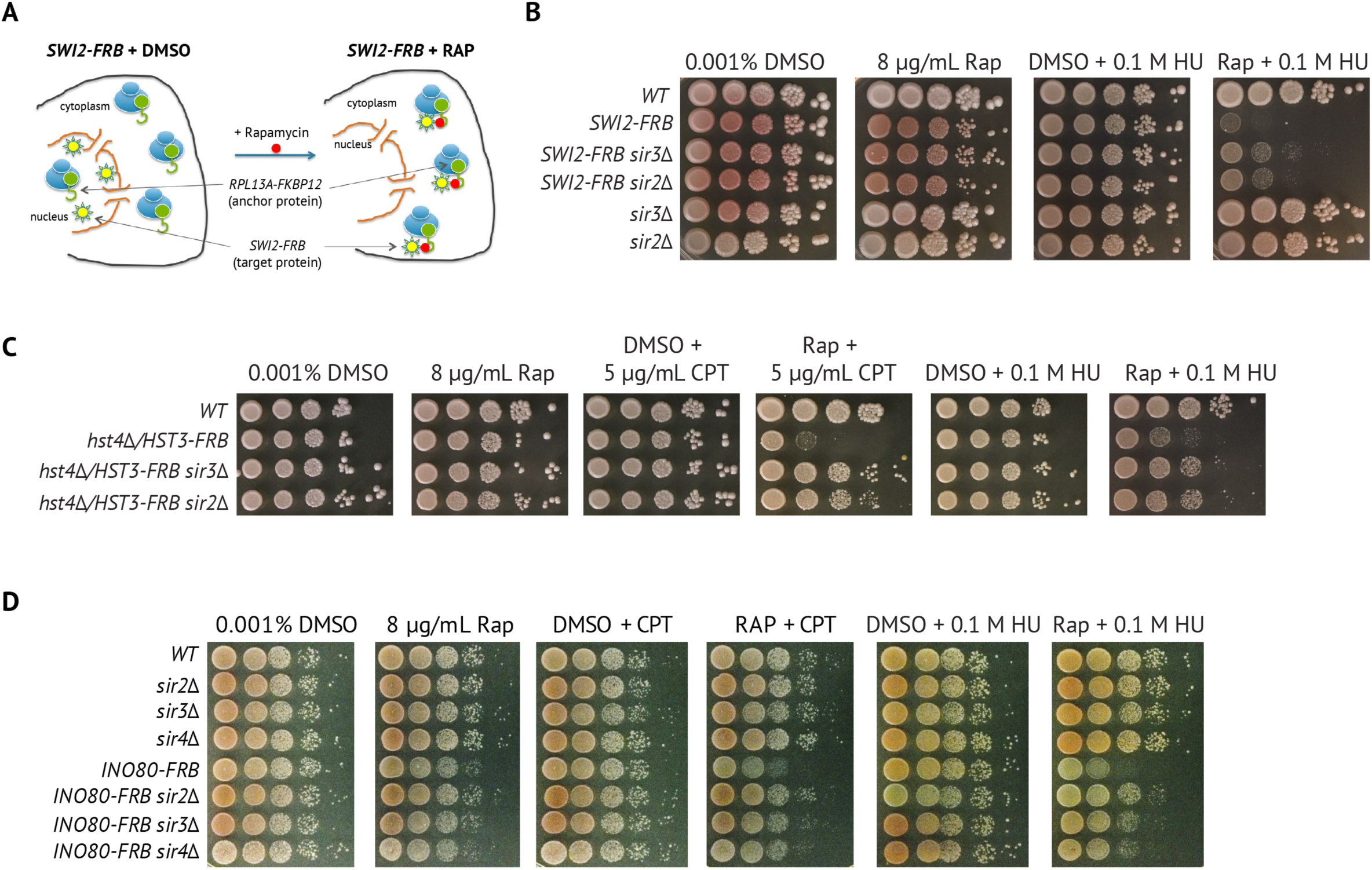
Absence of Sir2 *or* Sir3 partially suppresses phenotypes due to depletion of SWI/SNF, Hst3/Hst4, or INO80C. A) Schematic of the Anchor-away system to induce conditional depletion of nuclear proteins. Strains contain C-terminally tagged versions of the nucleo-cytoplasmic shuttling protein (*RPL13A-FKBP12;* green hook) and the *SWI2* gene locus (*SWI2-FRB;* yellow star) (left panel). Addition of Rapamycin (red dot) facilitates formation of a ternary complex between FKBP12 and FRB, rapidly depleting *SWI2-FRB* from the nucleus (right panel). B) Wild type, *sir2Δ*, and *sir3*Δ strains with or without the *SWI2*-FRB tag were spotted on 2% glucose media containing either DMSO solvent, 8µg/ml rapamycin (RAP) in the presence or absence of 0.1 M hydroxyurea (HU) and then grown for 3 days at 30°C. C) Spot assays as in B) for *hst4Δ/HST3-FRB* in the presence of or absence 0.1 M HU and 5 µg/mL Camptothecin (CPT). D) Spot assays as in B) for *INO80-FRB* in the presence or absence of 0.1 M HU and 5 µg/mL CPT.

We first compared growth rates of *SWI2*-FRB strains with or without the *SIR3* gene using spot assays. In the presence of DMSO solvent, growth rates of all strains are identical on rich media (Figure 2B), indicating that the *SWI2-FRB* fusion itself does not impair Swi2 function. In the presence of rapamycin, *SWI2-FRB* strains show a decrease in growth rate compared to the WT, consistent with nuclear depletion of Swi2 (Figure 2B). However, the *SWI2-FRB* strains have a milder growth defect compared to the *swi2Δ* (null) mutant (Figure 1), perhaps due to residual Swi2 present in the nucleus. Similar to the *swi2*Δ, depletion of *SWI2* also causes HU sensitivity, and this phenotype is partially suppressed by deletion of *SIR3*, suggesting an important link between Swi2 and Sir3 during replication stress (Figure 2B). Unlike the case with a deletion allele of *SWI2*, deletion of *SIR2* also partially suppressed the sensitivity of the SWI2-FRB strain (+Rap) to HU, but to a lesser extent than deletion of *SIR3* (Figure 2B).

Previous work has shown that *SWI2* is required for transcriptional activation of the <u>r</u>ibo<u>n</u>ucleotide <u>r</u>eductase (RNR) genes in the presence of HU (Sharma *et al*. 2003). Consistent with this, we see a large reduction of these transcripts in the *swi2*Δ (Figure S1E, F). However, unlike the rescue of growth, the lower levels of RNR transcripts was not restored by the *sir3Δ* following depletion of Swi2. This observation suggests that in HU stress, SWI/SNF may antagonize Sir3 independent of transcription, possibly by assisting replication within SIR heterochromatin.

### Deletion of *SIR2* or *SIR3* suppresses growth defects of *ino80* and *hst3/hst4* mutants

To investigate whether the suppression of growth defects by loss of Sir proteins is unique to SWI/SNF or a more common feature among chromatin modifying enzymes, we determined if deletion of *SIR2* and *SIR3* suppresses the growth defects caused by nuclear depletion of the histone H3 deacetylases, Hst3 and Hst4 (*Hst4Δ/HST3-FRB*), or the chromatin remodeler, INO80C (*INO80-FRB*). In the presence of rapamycin, both the *Hst4Δ/HST3-FRB* and *INO80-FRB* strains grow similarly to WT, but they are sensitive to stress conditions, such as when media contains camptothecin (CPT) or HU (Figure 2C, D). Strikingly, deletion of either *SIR2* or *SIR3* suppresses the growth defects caused by rapamycin-dependent depletion of either Ino80 or Hst3/Hst4 in the presence of HU or CPT (Figure 2C, D). Similarly, deletion of *SIR3* also partially suppresses an *ino80Δ* strain (Figure S2), confirming the results observed with INO80 anchor away. Thus, in addition to *SWI2*, the *SIR* genes also show genetic interactions with *HST3/HST4*, and *INO80*.

### Genotoxic stress is partially suppressed by *MAT* heterozygosity

Our genetic studies suggest that SWI/SNF, INO80C, and Hst3/Hst4 antagonize Sir proteins during replication stress. SIR heterochromatin prevents expression of the silent mating type loci, and consequently, loss of Sir proteins leads to expression of diploid-specific genes and suppression of haploid-specific genes (Rine and Herskowitz 1987; Goutte and Johnson 1988; Herskowitz 1989; Dranginis 1990). This pseudo-diploid state, in which both the MATa and MATα genes are expressed, is termed the *MAT* heterozygotic state. *MAT* heterozygosity has been shown to suppress the DNA repair defects of *rad* mutants (Valencia-Burton *et al*. 2006). Indeed, both MAT heterozygosity and *sir* mutants downregulate NHEJ and preferentially use HR for DNA repair (Valencia-Burton *et al*. 2006). To determine whether loss of Sir proteins suppresses replication stress phenotypes in a direct manner or an indirect manner due to pseudo-diploid effects, we investigated the impact of *MAT* heterozygosity. We introduced an episomal copy of *MATα* into the *MAT***a** anchor away strains to generate *MATa*/*MAT*α haploids. Interestingly, MAT heterozygosity partially suppressed the sensitivity of *SWI2-FRB* (+ Rap) during HU stress (Figure 3A), though the suppression was less than what was observed by deletion of *SIR3* (Figure 2B). In contrast, MAT heterozygosity did not suppress sensitivity of SWI2-FRB (+Rap) to CPT (Figure 3A). The sensitivity of the *hst4Δ/HST3-FRB* and *INO80-FRB* mutants to CPT, and to a lesser extent HU, was also partially suppressed by MAT heterozygosity (Figure 3B, C). Taken together, the results indicate that the replication stress sensitivity of strains depleted for Swi2, Ino80, or Hst3/Hst4 can be partially suppressed by *MAT* heterozygosity, either by deletion of *SIR* genes or by expressing the opposite mating type in haploid strains. However, the extent of suppression by the pseudo-diploid state is generally less than what is observed for loss of Sir proteins.

**Figure 3:**
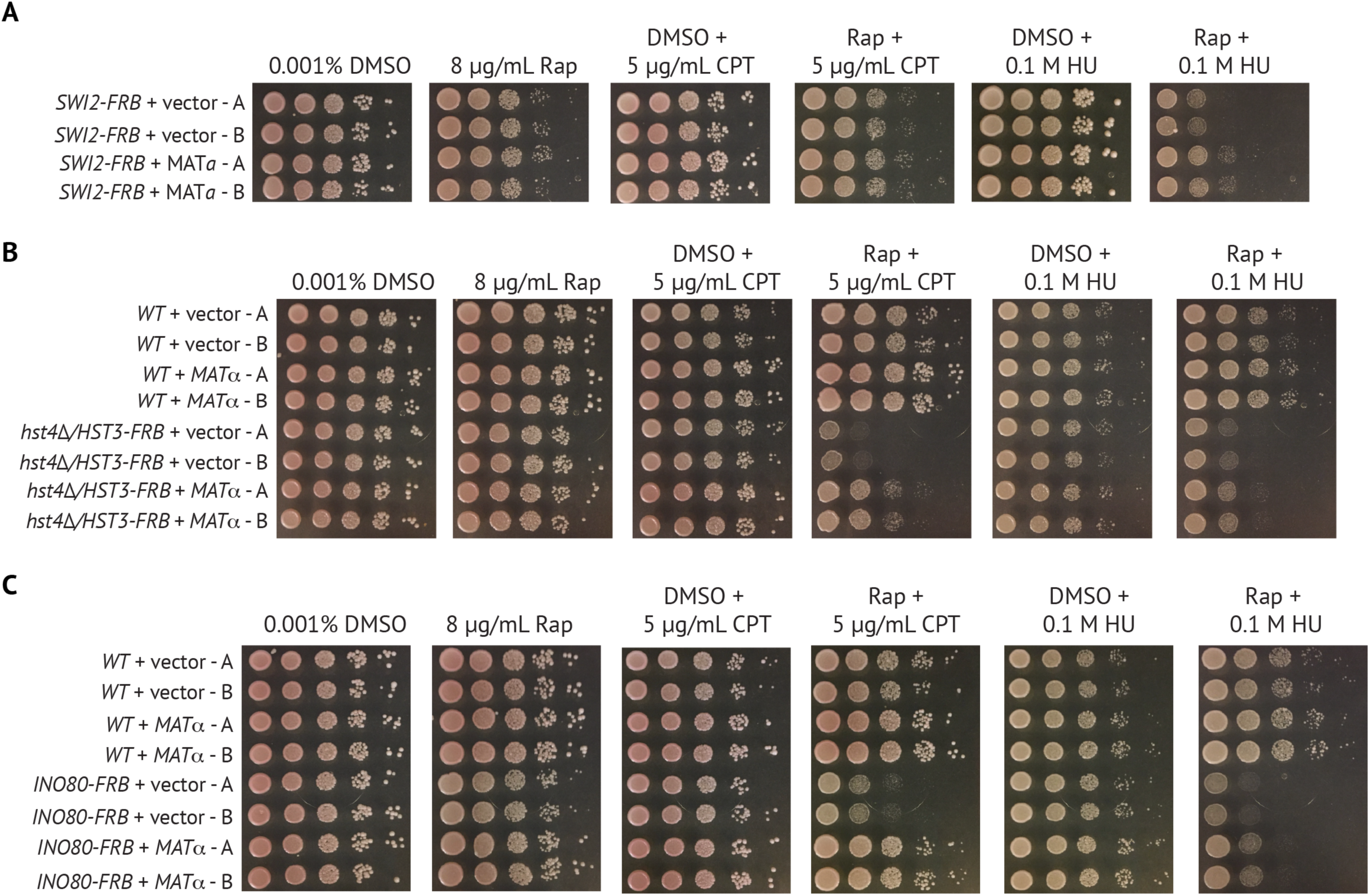
*MAT* heterozygosity partially suppresses replication stress phenotypes due to depletion of chromatin regulators. A) *SNF2*-FRB tag strains containing either empty vector or a vector expressing *MATa* were spotted on 2% glucose media containing either DMSO solvent, 8µg/ml rapamycin (RAP) in the presence or absence of 5 µg/mL Camptothecin (CPT) and 0.1 M HU, and then grown for 3 days at 30°C. Two transformants were tested. B) Spot assays as in A) for *hst4Δ/HST3-FRB* expressing *MATα* grown in the presence or absence of 5 µg/mL CPT and 0.1 M HU. C) Spot assays as in A) for *INO80-FRB* expressing *MATα* grown in the presence or absence of 5 µg/mL CPT and 0.1 M HU.

Expression of both MAT**a** and MATα alters the transcriptional profile by down regulating haploid-specific genes and upregulating diploid-specific genes (Herskowitz 1989). Thus, we sought to investigate which of these pathways are important for suppressing replication stress phenotypes. Previous work has shown that deletion of the haploid-specific gene *NEJ1*, required for non-homologous end joining (NHEJ), suppressed the growth defect of *rad55Δ* when exposed to DNA damage (Valencia-Burton *et al*. 2006). To determine whether loss of NHEJ also suppresses the phenotypes due to loss of chromatin modifiers, we deleted *NEJ1* in the anchor away strains.

Deletion of *NEJ1* did not suppress the growth defects of *SNF2-FRB*, *hst4Δ/HST3-FRB*, or *INO80-FRB* mutants when grown on rapamycin in the presence of HU or CPT (Figure S3A). We next tested the impact of two additional haploid-specific genes, *RME1* or *PST2*, which show genetic interactions with *RAD55* and *RAD51* during growth on CPT stress (Valencia-Burton *et al*. 2006). However, deletion of either *RME1* or *PST2* did not suppress the HU or CPT stress phenotypes of the *SWI2-FRB*, *hst4Δ/HST3-FRB*, or *INO80-FRB* (+Rap; Figure S3B, C). Thus, the underlying genetic basis for the *MAT*-dependent, partial suppression of stress phenotypes due to depletion of SWI/SNF, INO80, and Hst3/Hst4 remains unknown, though it does not appear to be due to loss of NHEJ or activation of meiotic transcriptional programs.

### Loss of Sir3 partially suppresses the transcriptional defects due to loss of SWI/SNF

Since loss of Sir3 has a large impact on phenotypes due to loss of Swi2 than *MAT* heterozygosity, we entertained the possibility that SWI/SNF may directly antagonize transcriptional repression by Sir3 at a subset of genes. To identify such transcriptional targets, we analyzed RNA profiles of isogenic wild type, *swi2Δ*, *sir3Δ,* and *swi2Δ sir3Δ* strains for 5716 ORFs using DNA microarrays. Consistent with published data, we observed that deletion of *SIR3* mis-regulates genes in the mating type cascade, with almost no other changes (Figure 4A *middle*) (Lenstra *et al*. 2011). In contrast, *SWI2* regulates 203 genes positively (FDR < 0.1 and LFC < −0.58) and 488 genes negatively (FDR < 0.1 and LFC > 0.58) (Figure 4A *top*). Many genes whose expression is known to be dependent on SWI/SNF, such as *SER3*, *YOR222W*, and the acid phosphatase genes, were altered as predicted (Figure 4B) (Sudarsanam *et al*. 2000). However, these SWI/SNF-dependent genes were unaffected by a deletion of *SIR3* (Figure 4B, third column).

**Figure 4:**
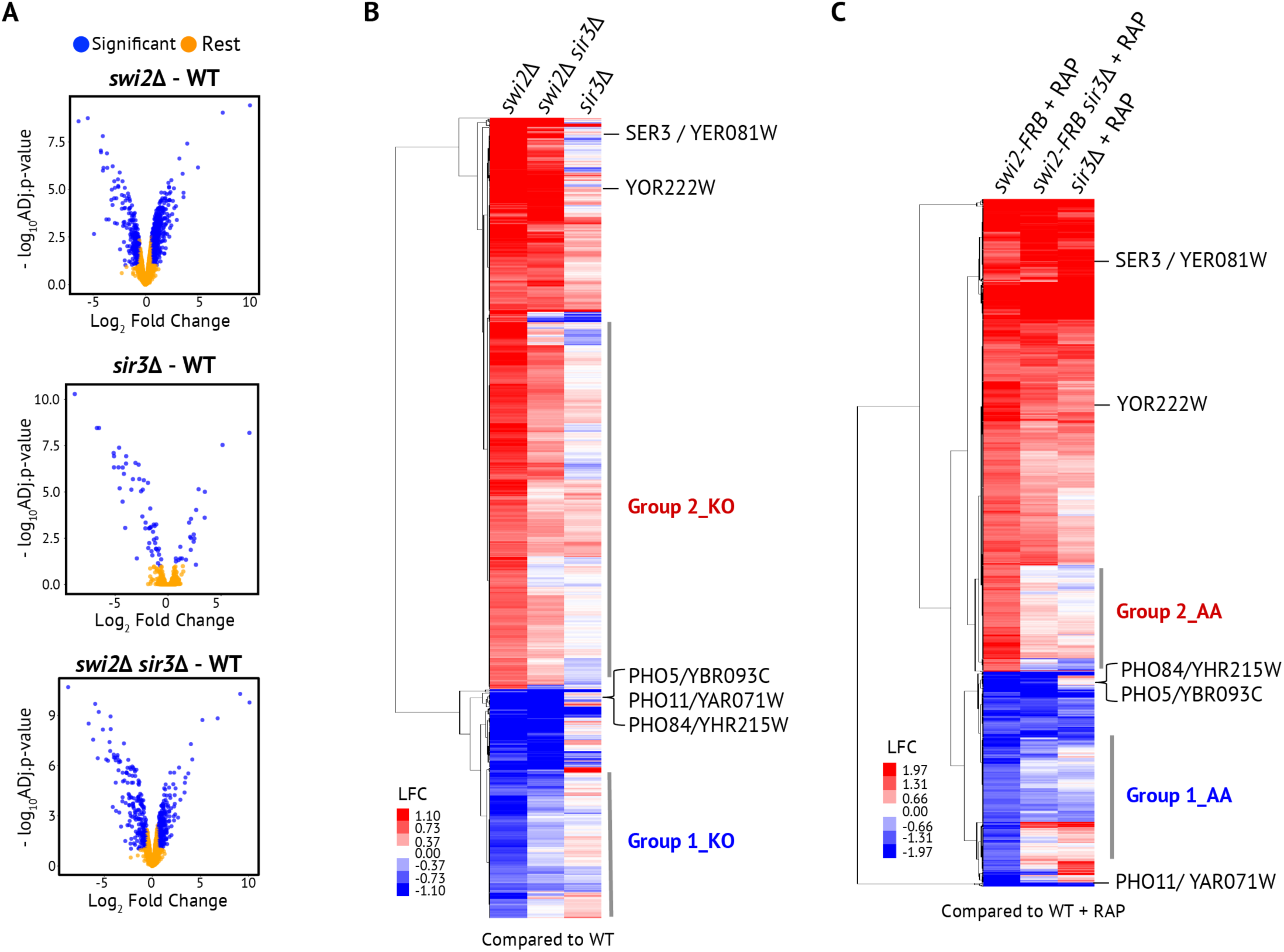
Whole-genome microarray analysis of *swi2*Δ and *SWI2*-AA strains. A) Volcano plots show transcripts that change significantly in the mutant compared to the wild type (WT) highlighted in blue (p.adj = FDR < 0.1 and Log2 Fold Change > 0.59). B) Heatmap of normalized RNA abundance for ORFs that are significantly down-regulated (n= 167) and up-regulated (n=488) in the *swi2Δ* arrays compared to WT. Corresponding values for these genes from *swi2Δ sir3Δ* and *sir3Δ* arrays compared to WT are also shown. Group 1_KO are defined as significantly down-regulated in the *swi2Δ* and comparatively de-repressed in *swi2Δ sir3Δ,* while Group 2_KO are defined as significantly up-regulated in the *swi2Δ* and comparatively reduced in *swi2Δ sir3Δ.* Examples of ORFs identified in previous studies that do not change in *swi2Δ sir3Δ* compared to *swi2Δ* (> ± 1.5 fold) are listed along the right. C) Heatmap of normalized RNA abundance for genes that are down-regulated (n=264) and up-regulated (n=193) in the *SWI2*-FRB compared to WT in the presence of 8µg/ ml of rapamycin (RAP). Corresponding values for these genes from *SWI2-FRB sir3Δ* and *sir3Δ* arrays compared to WT are also shown. ‘Group 1_AA’ and ‘Group 2_AA’ are defined essentially as described in B). Examples of ORFs identified in previous studies that do not change in *swi2Δ sir3Δ* compared to *swi2Δ* (> ± 1.5 fold) are listed along the right.

To identify genes that are regulated by both *SWI2* and *SIR3*, we first selected genes that changed significantly in the *swi2Δ* compared to wild type (Figure 4B), and we then performed hierarchical clustering and classified various sub-groups of interest. Genes that decrease significantly (LFC < −0.58 and FDR < 0.1) in *swi2*Δ and are restored to nearly wild type levels in the *swi2*Δ *sir3*Δ are defined as Group 1_KO (Table S1). The top gene ontology (GO) term category enriched in Group 1_KO is ribosome biogenesis/ ribosomal protein coding genes. This suggests that these genes require SWI/SNF to antagonize Sir3 to promote transcription. Indeed, prior studies have reported Sir3 binding to many ribosomal protein genes, using a *GAL-SIR3* inducible strain (Radman-Livaja *et al*. 2011). However, mRNA abundance of genes involved in ribosome biogenesis/ ribosomal proteins strongly anti-correlates with cellular growth rate and may confound our results (Airoldi *et al*. 2009).

To circumvent potential issues due to growth defects, we also analyzed RNA profiles from the anchor away strains. Transcriptional profiling following Swi2 depletion also identified many previously known SWI/SNF-dependent genes, including *YOR222W, SER3,* and the acid phosphatase genes (Figure 4C). Notably, genes involved in ribosome biogenesis were not identified in the anchor away datasets, consistent with the possibility that expression of these genes are linked to growth rates. To identify genes that might be co-regulated by Swi2 and Sir3, we again selected genes that changed by 1.5 fold or more after depletion of *SWI2-FRB*, and performed hierarchical clustering to identify subsets that are co-regulated by *sir3Δ*. Genes that *decrease* (LFC < −0.58) in *SWI2*-FRB and were restored to nearly wild type levels in the *SWI2*-FRB *sir3*Δ were defined as Group 1_AA (Figure 4C; Table S2). The top GO term category enriched in Group 1_AA is ion/ carbohydrate transport and primarily reflects the metabolic defects of *SWI2* mutants in carbon source utilization. The overlap between the Group1_AA and Group1_KO sets revealed a very select set of 28 genes that decrease following loss of SWI/SNF but are restored to nearly wildtype levels by inactivation of Sir3 (p-value of 8.7 x 10^-9^; Figure 5C; Hypergeometric test). This common subset of genes, consolidated as Group 1, corresponds to GO term categories of ‘cell cycle’, ‘cytokinesis’, and ‘lipid metabolism’ (Table S3). Notably, this set is enriched for genes expressed at the end of mitosis, which we previously showed to be SWI/SNF-dependent (Krebs, *et al*. 2000).

**Figure 5:**
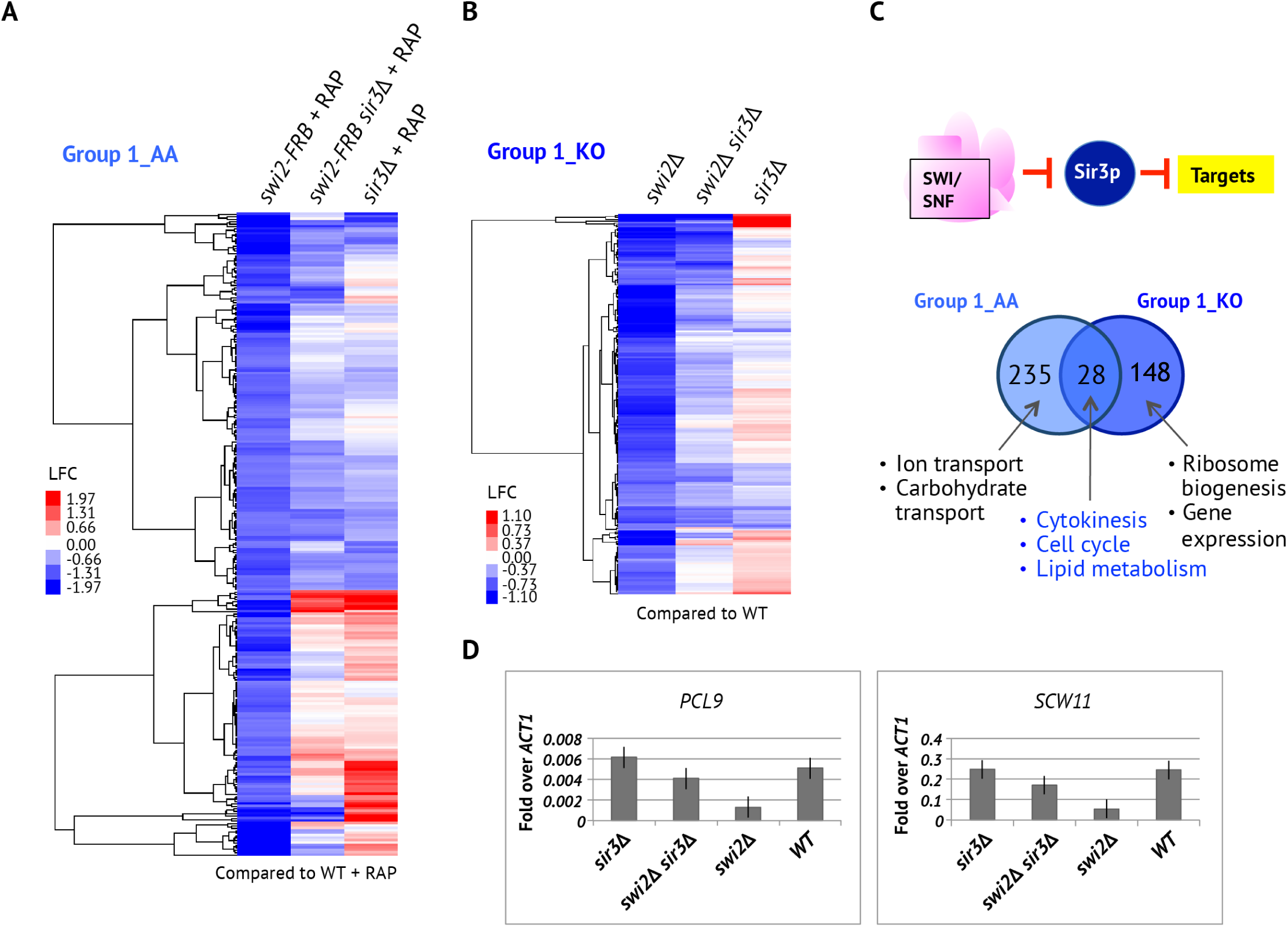
M/G1 expressed genes are regulated by SWI/SNF in a Sir3 dependent manner in both the *SWI2* anchor-away and *swi2Δ* strains. A) Heatmap of normalized RNA abundance for Group1_AA ORFs (n= 263) in the *SWI2-FRB*, *SWI2-FRB sir3Δ* and *sir3Δ* arrays compared to WT in the presence of 8µg/ ml of rapamycin (RAP) after hierarchical clustering. B) Heatmap of normalized RNA abundance for Group1_KO ORFs (n= 176) in the *swi2Δ*, *swi2Δ sir3Δ* and *sir3Δ* arrays compared to WT after hierarchical clustering. C) Venn diagram depicting the overlap of genes from Group 1_AA and Group 1_KO. GO terms specific and common to the knockout (KO) and anchor-away (AA) datasets are shown. D) RT-qPCR analysis of select Group 1 genes identified from both the knockout and anchor-away datasets sets.

Consistent with previous analyses, we also identified genes whose expression was increased by either the deletion of *SWI2* or by Swi2 nuclear depletion (Figure S4). Expression of a subset of these genes was restored to nearly wildtype levels by inactivation of Sir3, and these gene sets were designated Group 2_KO (n= 488; LFC < −0.58 and FDR < 0.1) and Group 2_AA (n= 192; LFC < −0.58 and FDR < 0.1). (Figure 4B,C and S4A,B; Table S4 and S5). However, overlap of Group 2_KO and Group 2_AA datasets revealed only 11 common genes (Figure S4C), suggesting that the upregulation of genes by loss of SWI/SNF is primarily due to an indirect effect of slow growth (Holstege *et al*. 1998b; Sudarsanam *et al*. 2000).

### Analysis of Sir3 binding at Group 1 target genes

The genetic and transcriptome analyses suggest that SWI/SNF antagonizes Sir3 to promote expression of specific genes. One prediction of this model is that Sir3 may accumulate at such target genes in the absence of SWI/SNF. To test the model, we analyzed Sir3 recruitment by chromatin immunoprecipitation (ChIP) in WT and *swi2*Δ mutants arrested in nocodazole. Nocodazole is a microtubule depolymerizing agent that blocks entry into mitosis and thus, cells accumulate at the G2/M border (Jacobs *et al*. 1988). Sir3 binding was measured using a native antibody to Sir3, as well as an anti-FLAG antibody in a strain expressing a *SIR3*-FLAG fusion from its endogenous locus. In both cases, ChIP analyses in the wild type strain demonstrated enrichment for Sir3 at the heterochromatic loci, HMR and TELVI-R. In the absence of Swi2, the occupancy of Sir3 is reduced at telomeres, consistent with a redistribution of Sir3 to ectopic loci (Figure S5). However, we did not observe significant changes in Sir3 enrichment at selected euchromatic target genes (Figure S5B).

## DISCUSSION

Establishing a separation between euchromatin and heterochromatin domains is crucial for cell function. The mechanisms that might actively exclude heterochromatin proteins from euchromatin domains remain poorly understood. Previously, we found that the SWI/SNF complex can remove the Sir3 heterochromatin protein from chromatin fibers *in vitro*, and here, we report genetic evidence that supports a role for SWI/SNF in disrupting the ability of Sir heterochromatin proteins to repress euchromatic gene expression. In particular, this activity of SWI/SNF appears crucial for proper expression of genes expressed at the end of mitosis.

### Genetic interactions between chromatin modifiers and *SIR* genes

Our *in vitro* studies indicated that the SWI/SNF chromatin remodeling enzyme was uniquely able to evict the Sir3 heterochromatin protein from chromatin fibers (Manning and Peterson 2014; Sinha *et al*. 2009). Here, we found that deletion of either *SIR3* or *SIR4* alleviated the growth defects of *swi2Δ* on media containing glucose, ethanol, or HU, but *sir3*Δ and *sir4*Δ did not significantly rescue the severe growth defects of *swi2*Δ on alternative carbon sources, such as raffinose or galactose. Likewise, inactivation of Sir proteins also alleviated the growth phenotypes of cells that were depleted of Swi2. Interestingly, the genetic interactions between genes encoding Sir proteins and SWI/SNF were not unique to this particular chromatin remodeler. Deletion of *SIR2* and *SIR3* also suppressed the growth defects of *ino80* and *hst3/hst4* mutants under genotoxic stress (Figure 2C, D), indicating a potential common mechanism for suppressing replication stress.

How does loss of Sir proteins alleviate the phenotypes of mutants that lack chromatin modifying enzymes? One possibility is that suppression is due to indirect effects caused by the pseudo-diploid state of *sir* mutant cells. Deletion of *SIR* genes induces the expression the a1-α2 repressor, generating pseudo-diploid cells. Such haploid cells expressing both *MAT* genes show greater resistance to radiation and are more recombination proficient than cells expressing only *MATα* or *MATa* (Heude and Fabre 1993). We found that pseudo-diploid cells partially suppressed the phenotypes of the *INO80-FRB* and *hst4Δ/HST3-FRB* strains to a similar level as deletion of *SIR* genes, especially for growth on CPT (Figure 3B,C), indicating the some genetic interactions between INO80C and Hst3/Hst4 with Sir proteins may be largely indirect. In contrast, although the pseudodiploid state partially suppressed the HU stress phenotype of the *SNF2-FRB* strain (Figure 3A), the suppression was much less than that observed after deletion of *SIR3* (Figure 2B). This suggests there exists both a direct genetic interaction between SWI/SNF and Sir3, consistent with the ability of SWI/SNF to evict Sir3 *in vitro* (Manning and Peterson 2014), as well as an indirect interaction caused by *MAT* heterozygosity.

### Transcriptional profiling of *swi2Δ* and *swi2Δ sir3Δ* strains

Our transcriptional profiling data suggest that there may be at least two classes of SWI/SNF-dependent genes – those where SWI/SNF antagonizes Sir proteins, and a second group of genes that may require more “canonical” nucleosome remodeling activities. Our transcriptional profiling results are consistent with this view, as we identified many SWI/SNF-dependent genes whose transcriptional defect was not restored by loss of Sir3 and a separate set of genes where SWI/SNF appears to antagonize Sir3. Furthermore, we recently identified and characterized a separation of function allele of *SWI2* (*swi2Δ-Δ10R*) that generates a SWI/SNF complex that has normal levels of nucleosome remodeling activity but lacks the ability to evict Sir3 from chromatin fibers. (Manning and Peterson 2014).

For genes where SWI/SNF appears to antagonize Sir3, our microarray analyses revealed two categories of genes – the first group includes ribosomal biogenesis and ribosomal protein coding genes and the second include genes involved in cytokinesis and cell division. Notably, the first category of genes are likely to be sensitive to growth rate and thus changes in their expression are most likely due to indirect effects (Airoldi *et al*. 2009) (Figure 5C). In contrast, genes involved in mitotic exit were identified in RNA profiles from both gene deletion strains and from the anchor away system. This suggest that defects in expression of these genes are likely to be independent of the growth defects of the swi2 deletion strain and may be direct targets where SWI/SNF antagonizes Sir3. These data lend mechanistic insight to previous findings that SWI/SNF promotes expression of genes involved in mitotic exit and that *SWI/SNF* mutants are defective in exiting mitosis (Krebs *et al*. 2000).

### Genes expressed during mitosis are dependent on SWI/SNF to antagonize Sir3

Why might Sir3 only impact genes that are expressed during mitosis? Cytological studies have shown that Sir3 localizes to discrete foci during the majority of the cell cycle, reflecting its heterochromatic localization (Laroche *et al*. 2000). In contrast, Sir3 shows a diffuse, nuclear staining pattern during mitosis, consistent with more promiscuous binding to both euchromatic and heterochromatic sites. However, we do not observe increased Sir3 occupancy at selected gene promoter regions by ChIP qPCR analyses in G2/M arrested cells, compared to asynchronous cell populations (Figure S5B), though a diffuse localization of Sir3 may not be detectable by ChIP-qPCR. Likewise, we did not detect changes in euchromatic Sir3 occupancy in the absence of SWI/SNF, though Sir3 levels were decreased from telomeric regions, consistent with previous reports (Dror and Winston 2004; Manning and Peterson 2014) (Figure S5). Currently, we favor a model in which Sir3 delocalizes from heterochromatic sites during mitosis, leading to the binding of Sir3 to euchromatic regions, perhaps facilitated by the deacetylated state of transcribed gene coding regions. Sir3 may bind in a diffuse manner across euchromatic genes, limiting the detection of Sir3 by ChIP analyses. We envision that SWI/SNF action may be required to remove Sir3, facilitating expression of these cell cycle regulated genes. Notably, this role for SWI/SNF would be distinct from the typical nucleosome remodeling activities of SWI/SNF.

## ACKNOWLEDGEMENTS

We thank Phyllis Spatrick and the Genomics Core Facility at UMass Medical School for assistance with sample processing. The pJR156 and pJR157 *MAT* plasmids were a gift from Prof. Jasper Rine at the University of California, Berkeley. This work was supported by grants from the NIH to C.L.P. (R35 GM122519) and to J.L.F. (F32 GM119229).

---

**Figure S1:**
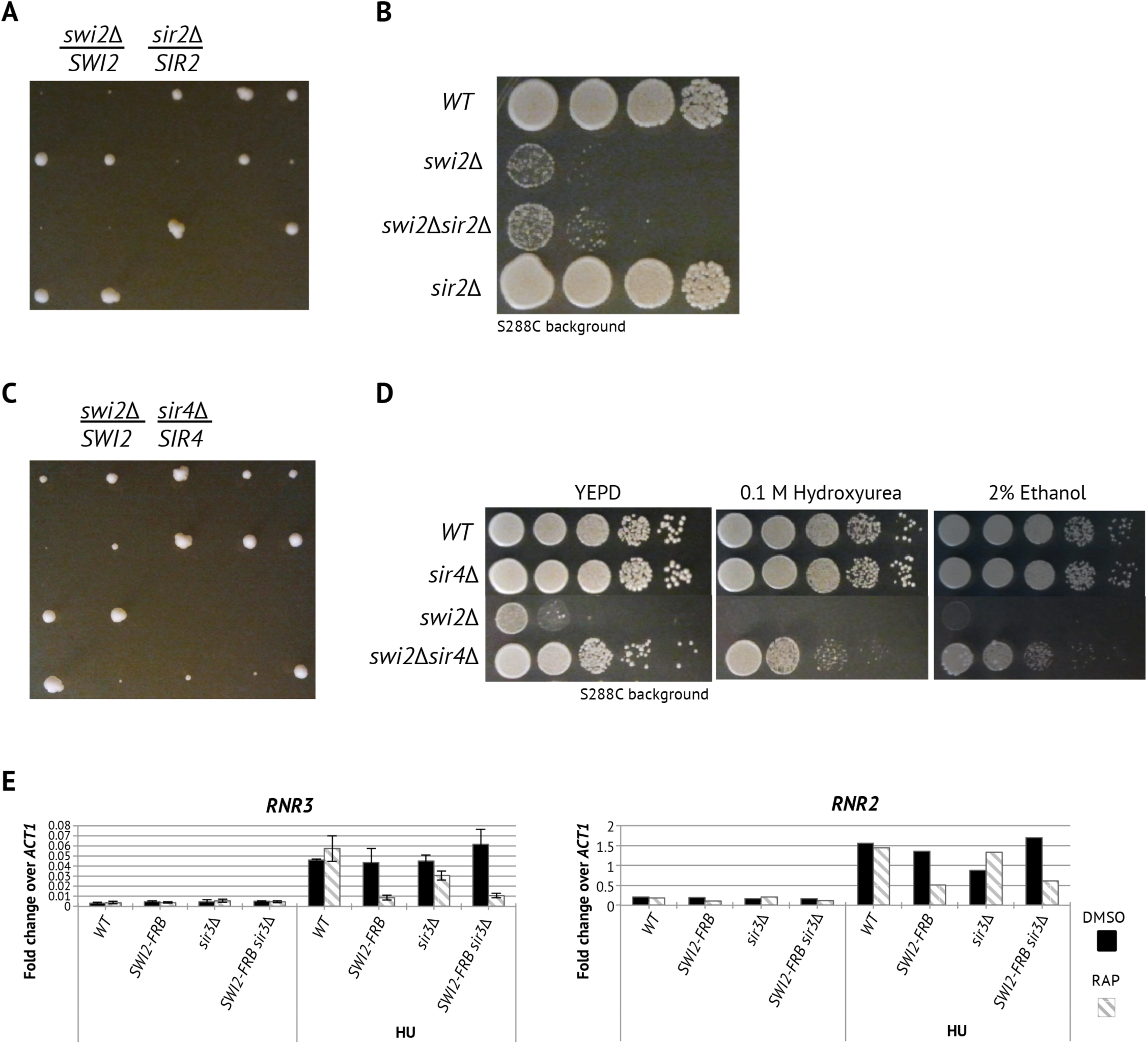
Gene expression and genetic interactions of *SIR3*, *SIR2,* and *SIR4* with *SWI2*. A, B) Absence of *SIR2* does not suppress growth defects of *swi2Δ*. C, D) Absence of *SIR4* suppresses the growth defects of *swi2Δ*. E) Absence of *SIR3* does not impact *RNR* gene expression and genetic interactions.

**Figure S2:**
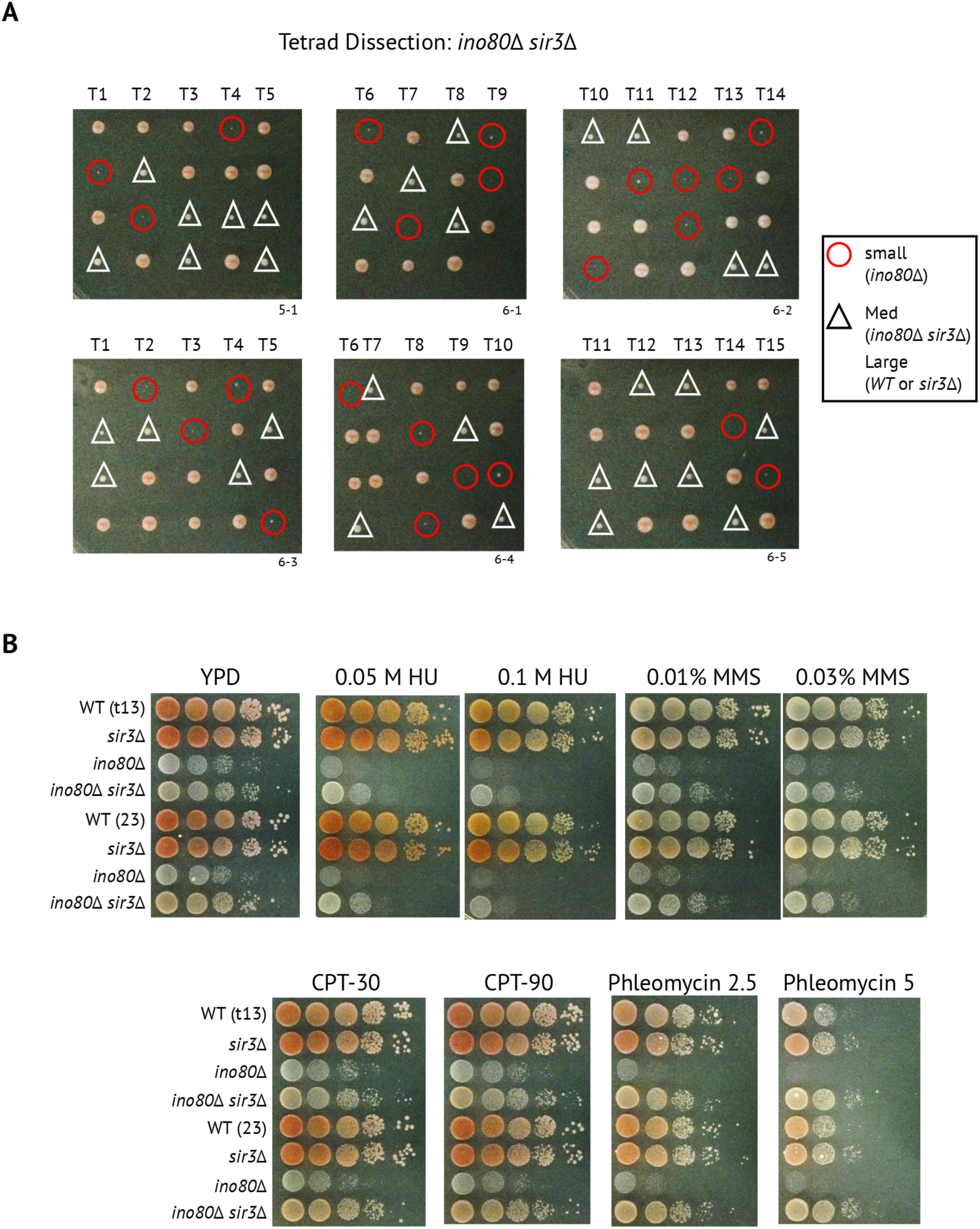
i*n*o80 growth defects are partially rescued by deletion of *SIR3*. A) Tetrad dissection plates of the *ino80Δ/INO80 sir3Δ/SIR3* heterozygous diploid on YEPD plates with the corresponding genotypes marked with symbols listed on the right. A single dissected spore yields an isogenic colony, imaged after 10 days. Relative size of each colony is representative of the growth rate. B) Spot assay on null mutants dissected from the W303 background. Equal cell numbers were spotted in consecutive ten-fold dilutions on agar plates with 2% glucose as the carbon source in the presence or absence of 0.05 M HU, 0.1 M HU, 0.01% Methyl methanesulfonate (MMS), 0.03% MMS, 30 µg/mL CPT, 90 µg/mL CPT, 2.5 µg/mL phleomycin, or 5 µg/mL phleomycin and imaged after 3 days.

**Figure S3:**
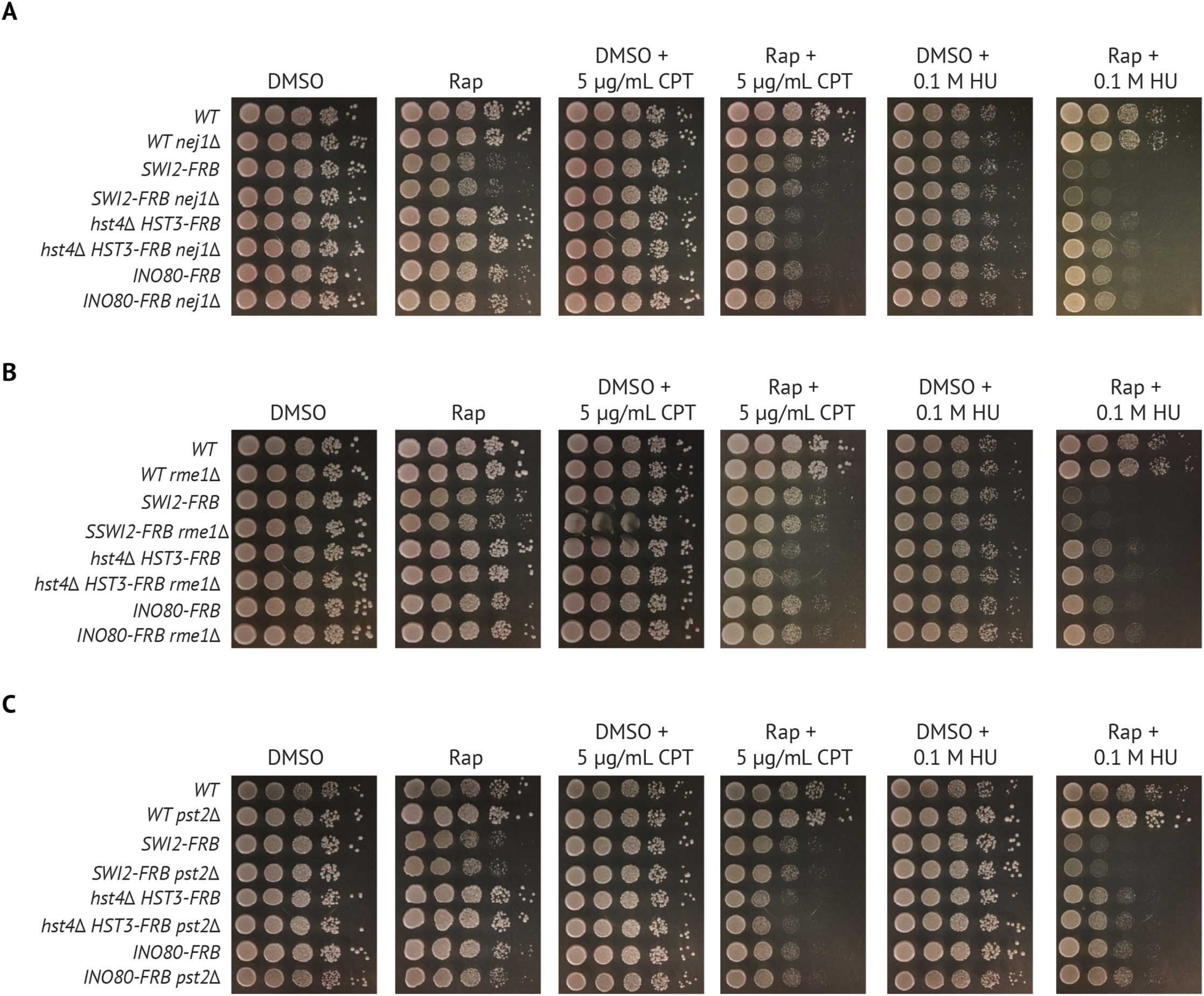
Growth defects due to depletion of chromatin regulators are not suppressed by deletion of haploid-specific genes *NEJ1*, *RME1*, or *PST2*. A) Wild type and *nej1Δ* strains with or without the *SNF2*-FRB, *hst4Δ/HST3-FRB*, or *INO80-FRB* were spotted (1/10 dilutions) on 2% glucose media containing either DMSO solvent, 8µg/ml rapamycin (RAP) in the presence or absence of 0.1 M HU and 5 µg/mL Camptothecin (CPT). Cells were then grown for 3 days at 30°C. B) Spot assays as in A) for *rme1Δ*. C) Spot assay as in A) for *pst2Δ*.

**Figure S4:**
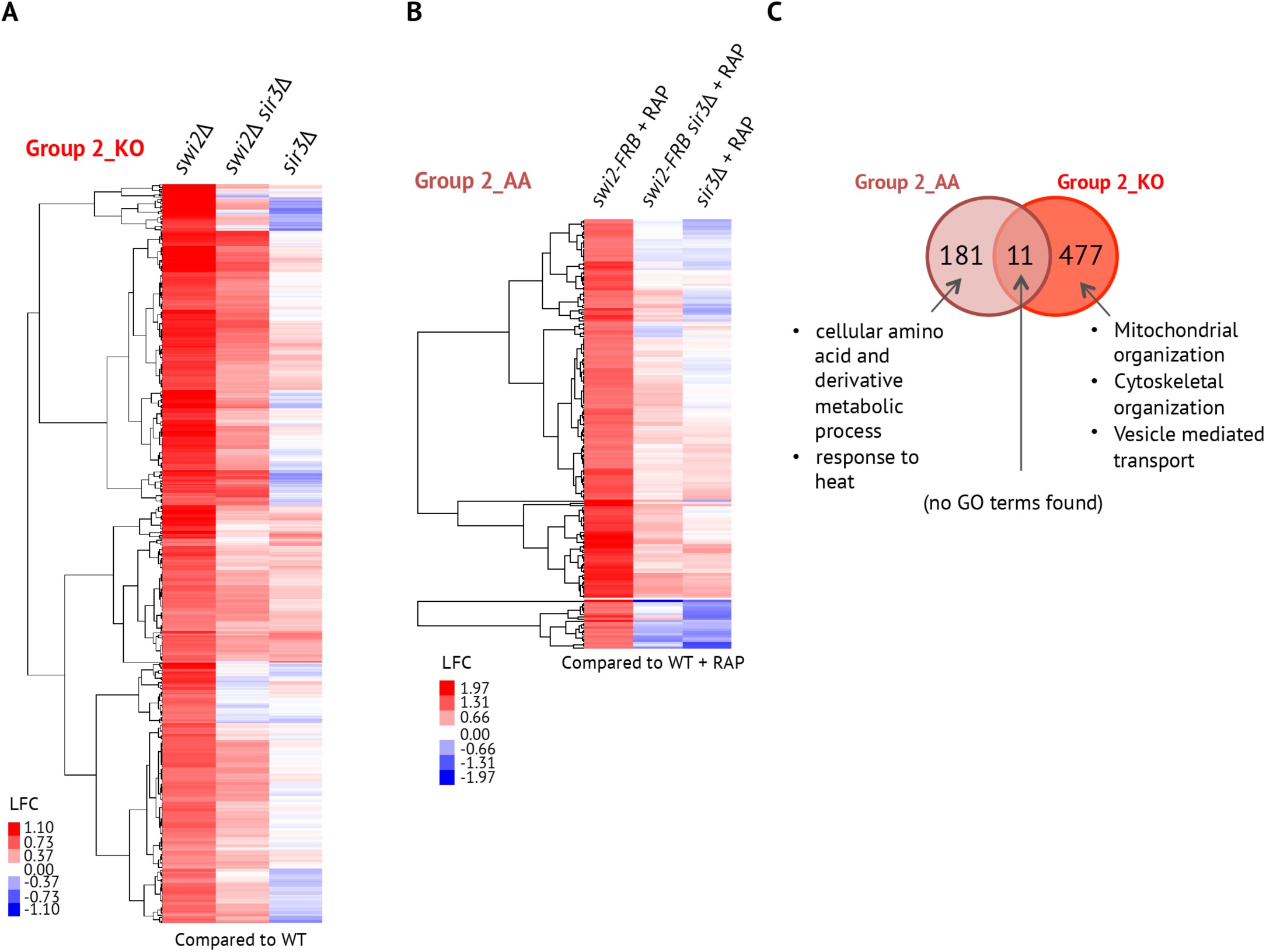
Overlap of Group 2 genes (those repressed by *SWI2*) between *swi2Δ* and *SWI2* **anchor-away strains**. A) Group 2_KO (n= 192) heatmap with strains compared to WT anchor away strain. B) Group 2_AA (n=488) heatmap with null mutants compared to WT. C) Overlap of the number of genes from Group 2_AA and Group 2_KO and the corresponding GO term categories.

**Figure S5:**
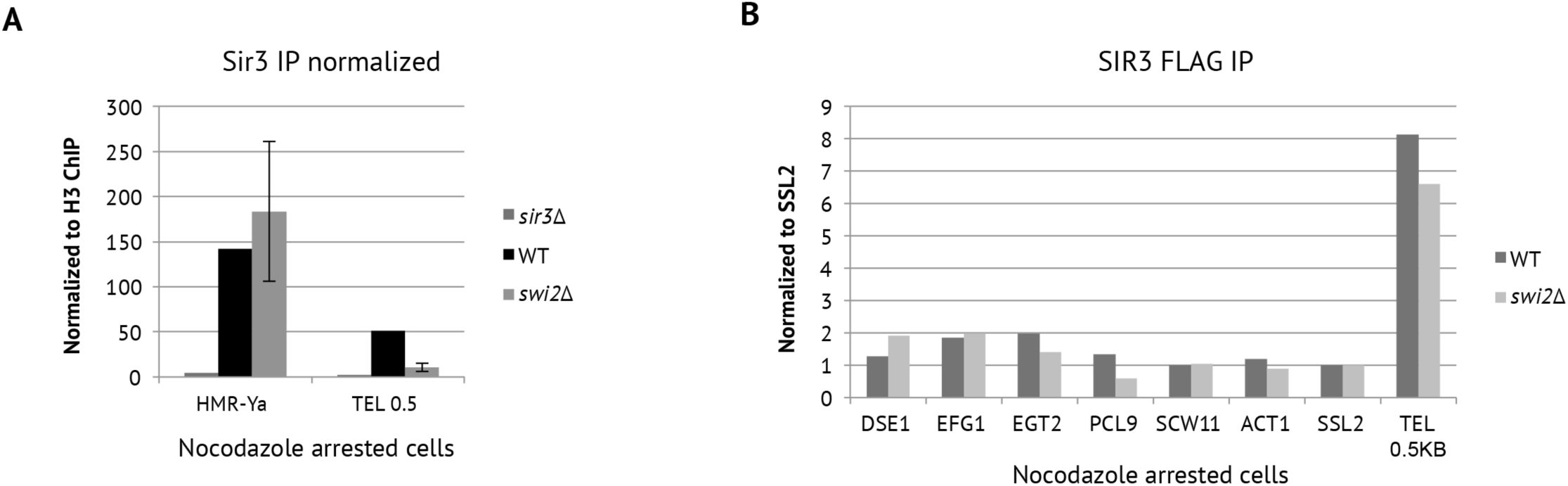
ChIP analysis of Sir3 occupancy. A) Chromatin immunoprecipitation (ChIP) for native Sir3 in nocodazole-arrested (G2/M boundary) cells at two heterochromatic loci in WT, *sir3*Δ and *swi2*Δ cells. B) ChIP for *SIR3*-FLAG in nocodazole-arrested (G2/M boundary) cells at promoters of *SWI2* dependent genes in WT and *swi2*Δ cells.

## SUPPLEMENTARY TABLES

**Table S1:** List of Group 1_KO genes. swi2D = swi2 null mutant; sir3D = sir3 null mutant, WT = Wildtype

| YORF | NAME | swi2D-WT | swi2Dsir3D-WT | sir3D-WT |
| --- | --- | --- | --- | --- |
| YOR107W | YOR107W | -1.736286 | -1.249234 | 0.802684 |
| YOR049C | YOR049C | -1.799513 | -1.528723 | 1.077021 |
| YOR032C | YOR032C | -1.190089 | -1.02068 | 1.545266 |
| YJL116C | YJL116C | -2.189504 | -0.596606 | 2.268662 |
| YIL169C | YIL169C | -0.686039 | -0.35075 | 2.457973 |
| YIL169C | YIL169C | -0.866575 | -0.786447 | 2.38699 |
| YOL155C | YOL155C | -0.771853 | -0.763125 | 2.132386 |
| YLL012W | YLL012W | -1.095636 | -0.664814 | 0.251892 |
| YMR182W-A | YMR182W | -1.19808 | -0.661026 | 0.069504 |
| YER072W | YER072W | -1.036205 | -0.931685 | -0.004959 |
| YBR238C | YBR238C | -1.113228 | -1.032084 | 0.050758 |
| YGR079W | YGR079W | -1.056979 | -0.935159 | 0.135865 |
| YNL141W | YNL141W | -1.263049 | -0.889395 | 0.046825 |
| YKL185W | YKL185W | -0.871451 | -0.845494 | -0.158825 |
| YHR049W | YHR049W | -0.982429 | -0.822014 | -0.31239 |
| YER062C | YER062C | -0.938457 | -0.793679 | -0.232057 |
| YOL002C | YOL002C | -0.883949 | -0.611531 | -0.186142 |
| YOR101W | YOR101W | -0.89683 | -0.624431 | -0.017271 |
| YLR008C | YLR008C | -0.812653 | -0.574562 | -0.044313 |
| YFL004W | YFL004W | -0.986432 | -0.698759 | 0.025827 |
| YBR085W | YBR085W | -1.020654 | -0.603666 | -0.067401 |
| YDR384C | YDR384C | -0.95195 | -0.607941 | 0.163866 |
| YJR060W | YJR060W | -0.792775 | -0.641355 | 0.115003 |
| YPR054W | YPR054W | -0.786343 | -0.572491 | 0.119735 |
| YLR300W | YLR300W | -0.593589 | -0.854639 | 0.03546 |
| YER044C | YER044C | -0.76323 | -0.995545 | 0.009447 |
| YPL092W | YPL092W | -0.784119 | -0.84839 | 0.077845 |
| YML075C | YML075C | -0.66478 | -0.742785 | 0.31812 |
| YPL066W | YPL066W | -0.538909 | -0.641721 | 0.178508 |
| YMR030W-A | YMR030W | -0.616896 | -0.454838 | 0.084452 |
| YJL117W | YJL117W | -0.600328 | -0.511951 | 0.060395 |
| YMR032W | YMR032W | -0.641548 | -0.584348 | 0.017399 |
| YER056C | YER056C | -0.658738 | -0.60332 | -0.057556 |
| YHR022C | YHR022C | -0.730672 | -0.465317 | 0.488107 |
| YNR009W | YNR009W | -0.639524 | -0.539942 | 0.276325 |
| YPL068C | YPL068C | -0.639863 | -0.390321 | 0.311581 |
| YDR044W | YDR044W | -0.787838 | -0.718158 | 0.576189 |
| YPL057C | YPL057C | -1.304416 | -0.453506 | -0.09573 |
| YNL327W | YNL327W | -1.419537 | -0.456658 | -0.035435 |
| YGR272C | YGR272C | -1.318424 | -0.351872 | -0.051133 |
| YJR048W | YJR048W | -1.257396 | -0.326339 | 0.014971 |
| YOL019W | YOL019W | -1.288769 | -0.387384 | 0.11906 |
| YKL099C | YKL099C | -1.150759 | -0.220279 | 0.07291 |
| YHR066W | YHR066W | -1.304707 | -0.185313 | 0.03815 |
| YOR095C | YOR095C | -0.997563 | -0.35532 | 0.050813 |
| YAL059W | YAL059W | -0.994499 | -0.398791 | 0.0139 |
| YNL162W-A | YNL162W- | -1.05393 | -0.313222 | -0.022577 |
| YIL158W | YIL158W | -1.038477 | -0.298129 | 0.012811 |
| YMR239C | YMR239C | -1.079968 | -0.323804 | 0.015705 |
| YDR184C | YDR184C | -1.117233 | -0.405462 | 0.066926 |
| YPL146C | YPL146C | -1.097313 | -0.341373 | 0.089128 |
| YLR073C | YLR073C | -1.124565 | -0.367879 | -0.154846 |
| YHR052W | YHR052W | -1.096203 | -0.388025 | -0.038756 |
| YMR015C | YMR015C | -1.192328 | -0.529732 | -0.121981 |
| YEL040W | YEL040W | -1.140605 | -0.537784 | -0.084598 |
| YNL024C | YNL024C | -1.124738 | -0.668689 | -0.176781 |
| YOR072W-B | YOR072W- | -1.179767 | -0.501843 | -0.313012 |
| YMR011W | YMR011W | -1.047188 | -0.494915 | -0.209743 |
| YDL179W | YDL179W | -1.4164 | -0.376551 | -0.21954 |
| YGR271C-A | YGR271C- | -1.367881 | -0.301543 | -0.243718 |
| YER124C | YER124C | -1.394713 | -0.365854 | -0.539364 |
| YPL061W | YPL061W | -1.293322 | -0.005912 | -0.610692 |
| YBL028C | YBL028C | -1.025202 | -0.206871 | -0.053283 |
| YGL028C | YGL028C | -0.98148 | -0.264637 | -0.118008 |
| YCR020W-B | YCR020W- | -1.018662 | -0.087625 | -0.049605 |
| YJL011C | YJL011C | -0.792154 | -0.010188 | -0.011872 |
| YPL158C | YPL158C | -0.814284 | 0.01076 | -0.092527 |
| YHR196W | YHR196W | -0.863989 | -0.165838 | -0.124069 |
| YLR287C | YLR287C | -0.782722 | -0.099197 | -0.197116 |
| YOL020W | YOL020W | -0.739805 | -0.203711 | -0.071481 |
| YOR159C | YOR159C | -0.673594 | -0.138845 | -0.077676 |
| YPL227C | YPL227C | -0.680673 | -0.181158 | -0.113845 |
| YLR264W | YLR264W | -0.649254 | -0.204616 | -0.20628 |
| YPL163C | YPL163C | -0.819237 | -0.39144 | -0.165403 |
| YCR043C | YCR043C | -0.828681 | -0.340857 | -0.196241 |
| YBL055C | YBL055C | -0.690262 | -0.333605 | -0.148308 |
| YKL164C | YKL164C | -0.886037 | -0.389714 | -0.091537 |
| YLR285C-A | YLR285C-A | -0.812289 | -0.440132 | -0.07874 |
| YKR061W | YKR061W | -0.855789 | -0.461683 | 0.016426 |
| YNL075W | YNL075W | -0.873769 | -0.303583 | 0.06103 |
| YOR119C | YOR119C | -0.88207 | -0.345121 | 0.01433 |
| YOR004W | YOR004W | -0.878897 | -0.356542 | 0.055945 |
| YGR081C | YGR081C | -0.886528 | -0.245544 | -0.064313 |
| YDL063C | YDL063C | -0.8893 | -0.297907 | -0.073846 |
| YML043C | YML043C | -0.904041 | -0.215257 | -0.006759 |
| YNL300W | YNL300W | -0.767885 | -0.357339 | 0.057677 |
| YPL165C | YPL165C | -0.804397 | -0.320086 | 0.053357 |
| YOL144W | YOL144W | -0.742002 | -0.28505 | 0.092757 |
| YML080W | YML080W | -0.808155 | -0.343047 | -0.05663 |
| YNL034W | YNL034W | -0.798485 | -0.305995 | -0.046402 |
| YNL119W | YNL119W | -0.738471 | -0.290834 | -0.06127 |
| YDR399W | YDR399W | -0.730769 | -0.433201 | -0.034015 |
| YDL042C | YDL042C | -0.852518 | -0.28447 | 0.312551 |
| YJL122W | YJL122W | -0.801952 | -0.352235 | 0.287684 |
| YGR041W | YGR041W | -0.81707 | -0.416311 | 0.221974 |
| YLR063W | YLR063W | -0.785169 | -0.351531 | 0.189522 |
| YHR169W | YHR169W | -0.921531 | -0.335304 | 0.213337 |
| YLR363W-A | YLR363W- | -0.940268 | -0.214857 | 0.080432 |
| YNL301C | YNL301C | -0.952251 | -0.208943 | 0.084877 |
| YDR299W | YDR299W | -0.920373 | -0.242432 | 0.116508 |
| YGL029W | YGL029W | -0.974169 | -0.244291 | 0.044468 |
| YHR197W | YHR197W | -0.983111 | -0.326906 | 0.130451 |
| YLL011W | YLL011W | -0.978058 | -0.249897 | 0.129031 |
| YNL182C | YNL182C | -1.00086 | -0.256279 | 0.160956 |
| YOR287C | YOR287C | -1.026403 | -0.257126 | 0.214451 |
| YOL080C | YOL080C | -1.069416 | -0.271948 | 0.147793 |
| YLR068W | YLR068W | -0.939414 | -0.062827 | 0.087384 |
| YOR342C | YOR342C | -0.809384 | -0.059979 | 0.125933 |
| YCR016W | YCR016W | -0.852698 | -0.095476 | 0.052606 |
| YOR264W | YOR264W | -0.85863 | 0.023682 | 0.160862 |
| YDR412W | YDR412W | -0.840776 | 0.039301 | 0.100001 |
| YPL245W | YPL245W | -0.926171 | 0.073276 | 0.256865 |
| YIL019W | YIL019W | -0.824764 | 0.091556 | 0.273614 |
| YDR083W | YDR083W | -0.915437 | -0.173763 | 0.247534 |
| YHR085W | YHR085W | -0.837462 | -0.253564 | 0.215562 |
| YDR324C | YDR324C | -0.829401 | -0.198156 | 0.256622 |
| YLR003C | YLR003C | -0.766628 | -0.120755 | 0.136361 |
| YGR187C | YGR187C | -0.740108 | -0.130188 | 0.16448 |
| YDR361C | YDR361C | -0.806214 | -0.157068 | 0.185872 |
| YPR137W | YPR137W | -0.771569 | -0.095819 | 0.253034 |
| YIL103W | YIL103W | -0.822611 | -0.177048 | -8.72E-04 |
| YJL148W | YJL148W | -0.804848 | -0.162139 | 0.02211 |
| YDR449C | YDR449C | -0.828149 | -0.23701 | 0.019955 |
| YIL091C | YIL091C | -0.829529 | -0.198132 | 0.05289 |
| YER127W | YER127W | -0.877939 | -0.199958 | 0.148394 |
| YNL111C | YNL111C | -0.809315 | -0.218915 | 0.120586 |
| YKR024C | YKR024C | -0.780613 | -0.240749 | 0.136802 |
| YDL043C | YDL043C | -0.748002 | -0.186336 | 0.044226 |
| YGR283C | YGR283C | -0.78912 | -0.186846 | 0.043388 |
| YLR347C | YLR347C | -0.71737 | -0.127022 | 0.065812 |
| YLR222C | YLR222C | -0.762311 | -0.144403 | 0.06022 |
| YFL034C-A | YFL034C-A | -0.621965 | -0.448212 | -0.283375 |
| YPR166C | YPR166C | -0.652454 | -0.411842 | -0.275068 |
| YPR106W | YPR106W | -0.62125 | -0.625154 | -0.222107 |
| YDR210W | YDR210W | -0.76796 | -0.353979 | -0.35536 |
| YJL096W | YJL096W | -0.585856 | -0.368384 | -0.041313 |
| YNL169C | YNL169C | -0.525778 | -0.303254 | -0.144911 |
| YPR108W-A | YPR108W- | -0.549497 | -0.331522 | -0.176694 |
| YDR070C | YDR070C | -0.790551 | -0.667047 | -0.574221 |
| YMR118C | YMR118C | -0.573588 | -0.551602 | -0.631359 |
| YML058W-A | YML058W- | -0.593191 | 0.265816 | -0.440319 |
| YML108W | YML108W | -0.495256 | 0.063711 | -0.148922 |
| YKL096W | YKL096W | -1.5604 | -0.110821 | 0.363457 |
| YMR230W-A | YMR230W | -1.686877 | -0.517905 | 0.255474 |
| YKL078W | YKL078W | -1.078741 | -0.409123 | 0.246475 |
| YDL241W | YDL241W | -1.03905 | -0.521772 | 0.4537 |
| YLR121C | YLR121C | -1.101156 | -0.093144 | 0.564482 |
| YHR094C | YHR094C | -1.207275 | 0.243913 | 0.274392 |
| YNR067C | YNR067C | -1.430648 | 0.257047 | 0.118234 |
| YDR222W | YDR222W | -1.421418 | 0.408224 | 0.079002 |
| YJR070C | YJR070C | -0.569932 | -0.180888 | 0.142184 |
| YLR409C | YLR409C | -0.618153 | -0.238689 | 0.185317 |
| YLR221C | YLR221C | -0.612013 | -0.171469 | 0.213431 |
| YOR340C | YOR340C | -0.615029 | -0.141354 | 0.225569 |
| YKL166C | YKL166C | -0.69773 | -0.1267 | 0.148319 |
| YJL069C | YJL069C | -0.678246 | -0.180436 | 0.173403 |
| YNL175C | YNL175C | -0.665957 | -0.110683 | 0.207458 |
| YLR099C | YLR099C | -0.403286 | -0.129096 | 0.199719 |
| YDR398W | YDR398W | -0.530144 | -0.079465 | 0.183036 |
| YPL069C | YPL069C | -0.607505 | 0.072517 | 0.178488 |
| YOL007C | YOL007C | -0.626065 | -0.02273 | 0.183157 |
| YOR091W | YOR091W | -0.6291 | -0.014712 | 0.163624 |
| YPR143W | YPR143W | -0.62508 | 0.010663 | 0.175941 |
| YDR087C | YDR087C | -0.654831 | -0.003588 | 0.136556 |
| YIL127C | YIL127C | -0.779948 | -0.014545 | 0.186459 |
| YDR021W | YDR021W | -0.740934 | 0.048523 | 0.200147 |
| YER082C | YER082C | -0.706368 | -0.012994 | 0.202725 |
| YOL010W | YOL010W | -0.638388 | 0.015929 | 0.237123 |
| YDR365C | YDR365C | -0.709428 | 0.021993 | 0.278121 |
| YAL025C | YAL025C | -0.635748 | -0.014059 | 0.325016 |
| YIL104C | YIL104C | -0.628021 | -0.131091 | 0.288274 |
| YBR247C | YBR247C | -0.694492 | -0.127991 | 0.286097 |
| YPL043W | YPL043W | -0.696074 | -0.090589 | 0.270935 |
| YBR267W | YBR267W | -0.668156 | -0.086142 | 0.323763 |
| YBR271W | YBR271W | -0.561659 | -0.021174 | 0.463252 |
| YER028C | YER028C | -0.63341 | 0.196135 | 0.762258 |

**Table S2:**
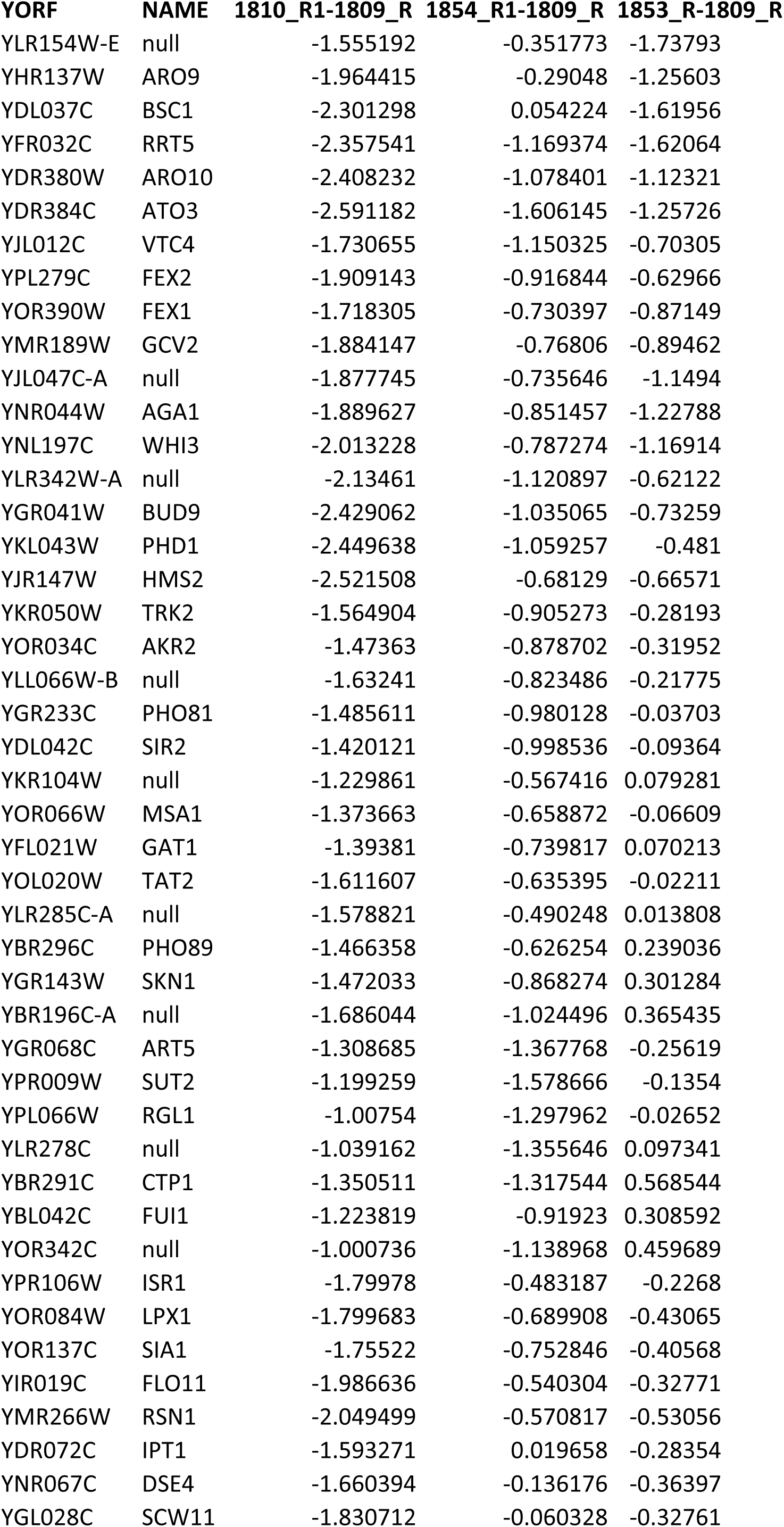
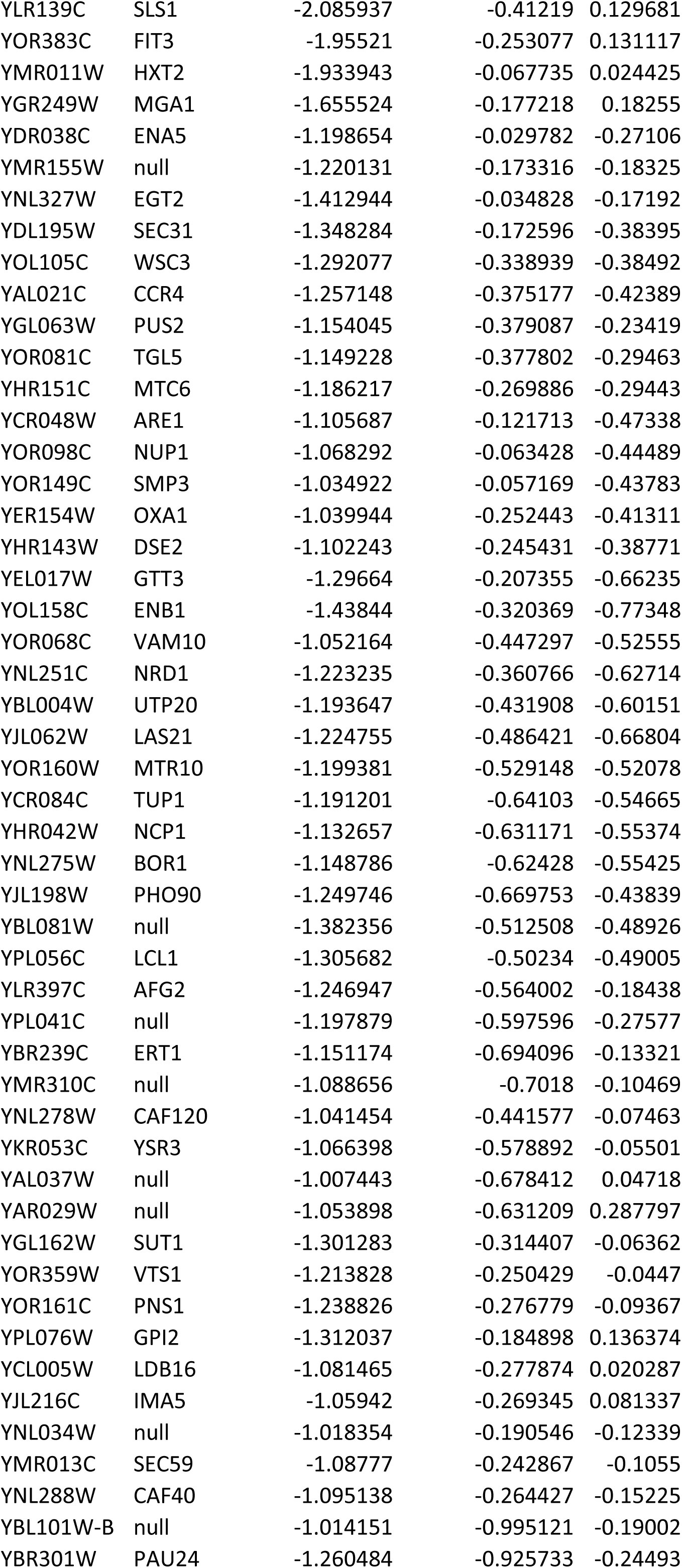
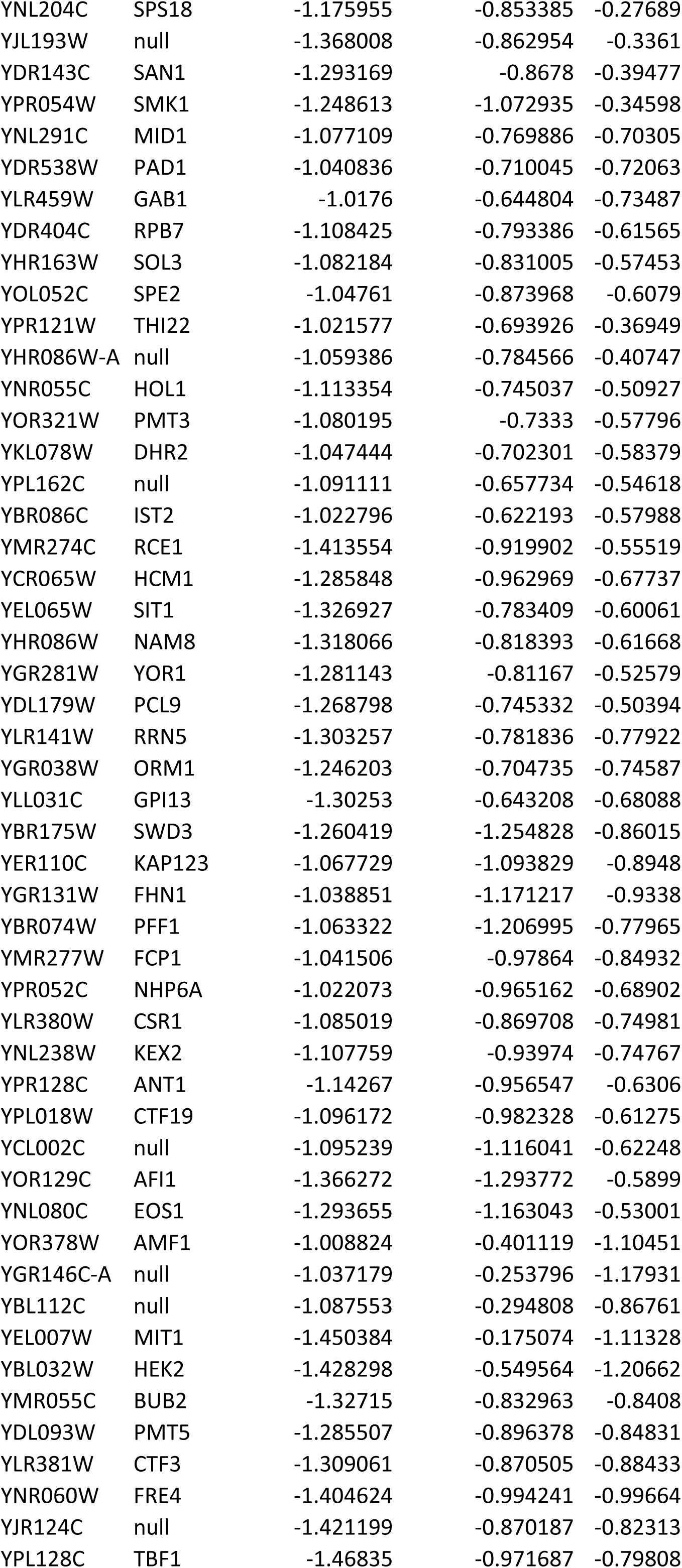
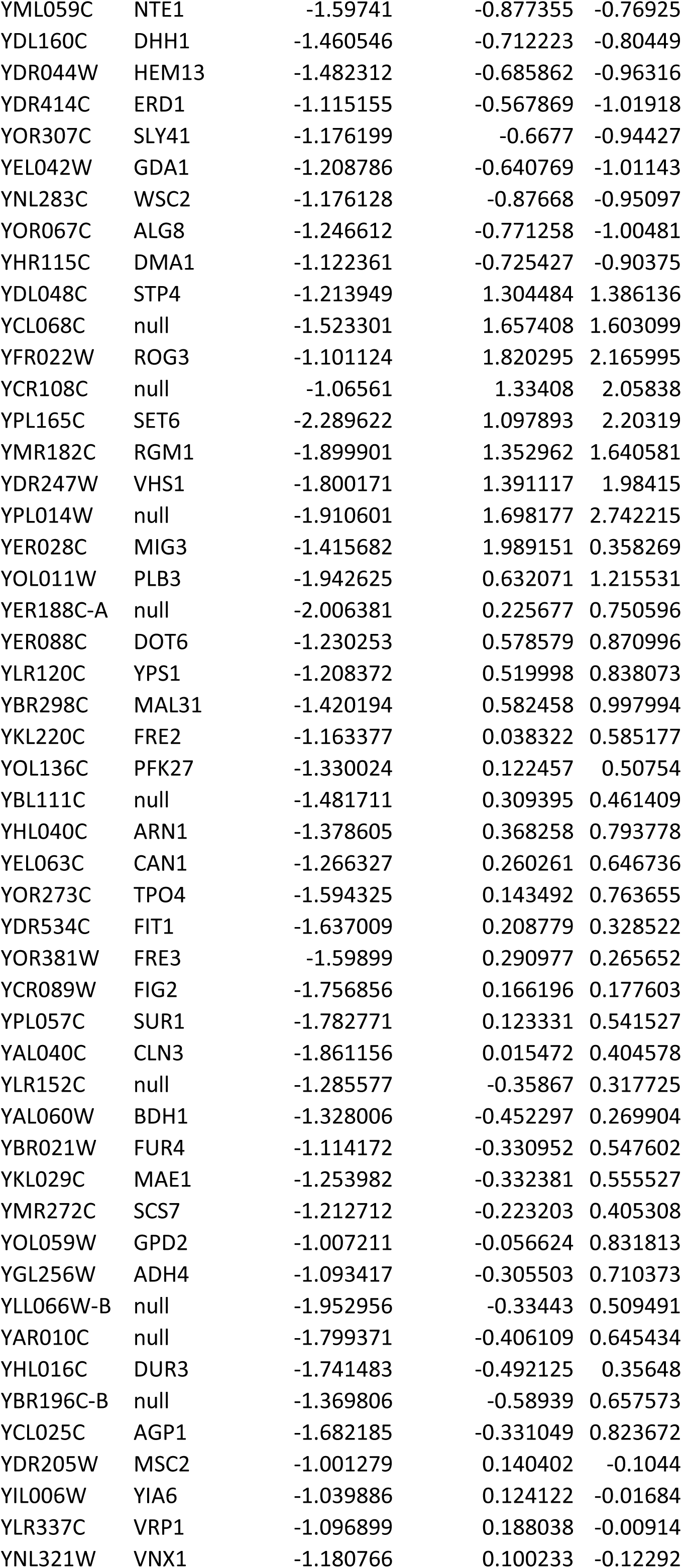
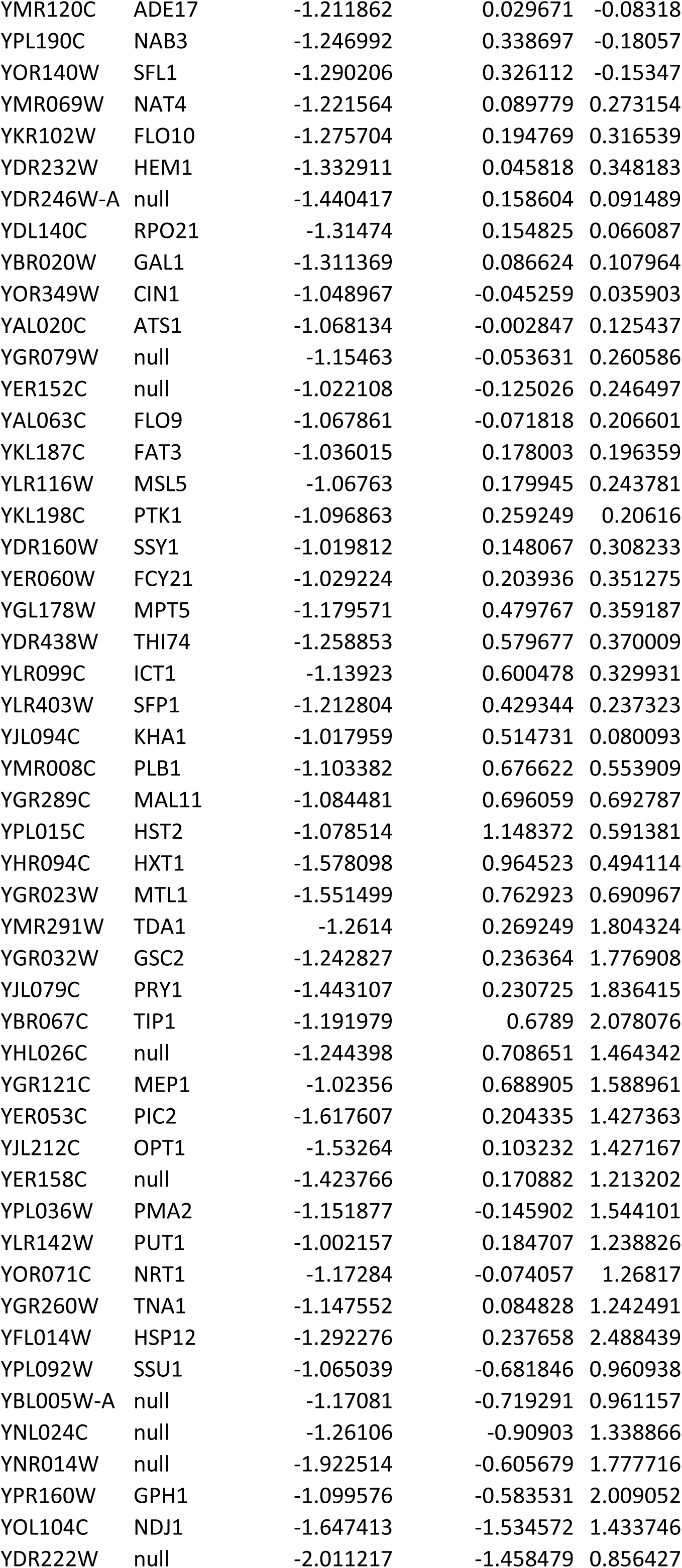
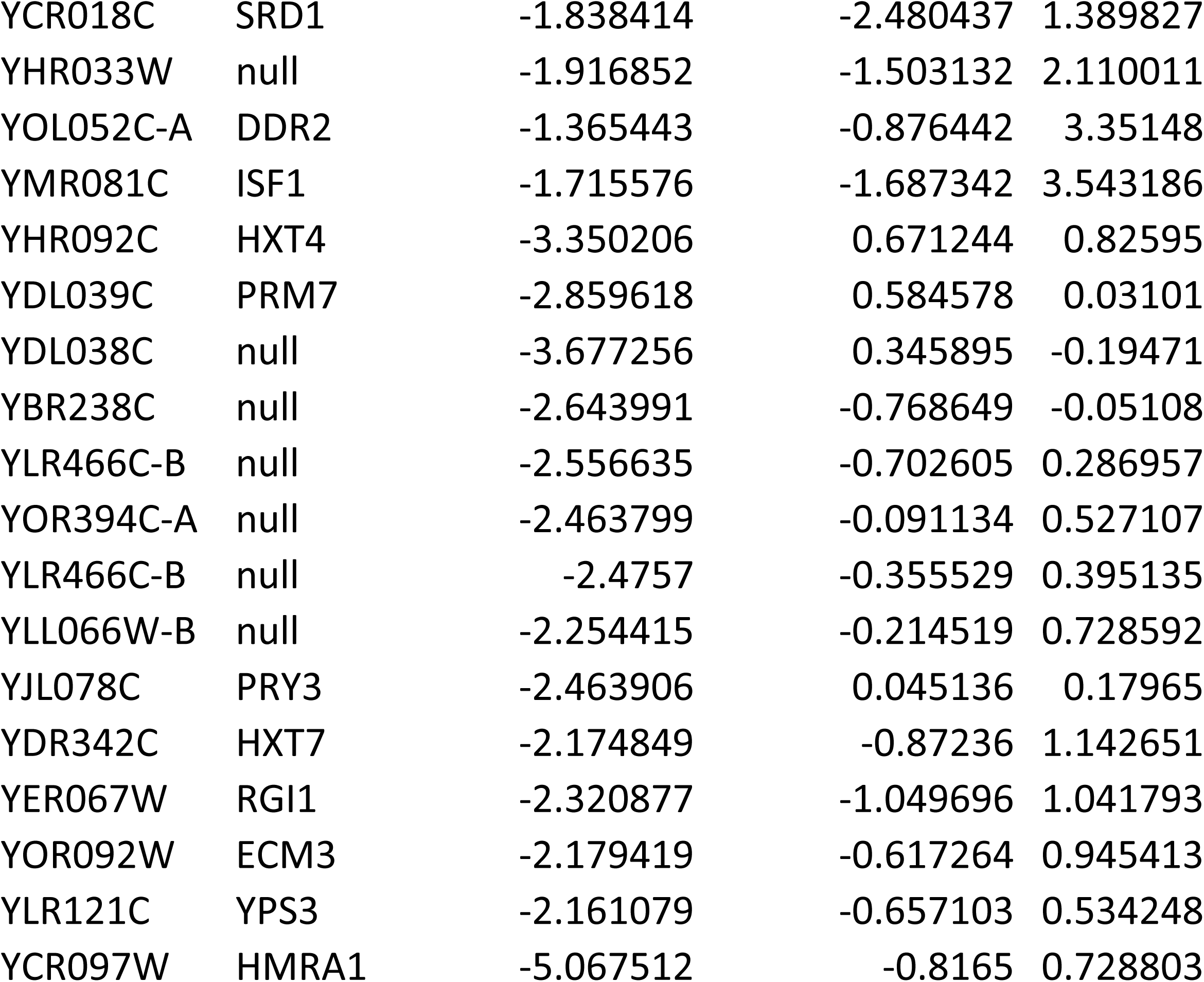
List of Group 1_AA genes. 1810 = SWI2 Anchor away; 1854 = SWI2 Anchor away sir3 null mutant; 1853 = sir3 null in Anchor away background; 1809 = Wildtype Anchor away background. _R = with Rapamycin

| YORF | NAME | 1810_R1-1809_R | 1854_R1-1809_R | 1853_R-1809_R |
| --- | --- | --- | --- | --- |
| YLR154W-E | null | -1.555192 | -0.351773 | -1.73793 |
| YHR137W | ARO9 | -1.964415 | -0.29048 | -1.25603 |
| YDL037C | BSC1 | -2.301298 | 0.054224 | -1.61956 |
| YFR032C | RRT5 | -2.357541 | -1.169374 | -1.62064 |
| YDR380W | ARO10 | -2.408232 | -1.078401 | -1.12321 |
| YDR384C | ATO3 | -2.591182 | -1.606145 | -1.25726 |
| YJL012C | VTC4 | -1.730655 | -1.150325 | -0.70305 |
| YPL279C | FEX2 | -1.909143 | -0.916844 | -0.62966 |
| YOR390W | FEX1 | -1.718305 | -0.730397 | -0.87149 |
| YMR189W | GCV2 | -1.884147 | -0.76806 | -0.89462 |
| YJL047C-A | null | -1.877745 | -0.735646 | -1.1494 |
| YNR044W | AGA1 | -1.889627 | -0.851457 | -1.22788 |
| YNL197C | WHI3 | -2.013228 | -0.787274 | -1.16914 |
| YLR342W-A | null | -2.13461 | -1.120897 | -0.62122 |
| YGR041W | BUD9 | -2.429062 | -1.035065 | -0.73259 |
| YKL043W | PHD1 | -2.449638 | -1.059257 | -0.481 |
| YJR147W | HMS2 | -2.521508 | -0.68129 | -0.66571 |
| YKR050W | TRK2 | -1.564904 | -0.905273 | -0.28193 |
| YOR034C | AKR2 | -1.47363 | -0.878702 | -0.31952 |
| YLL066W-B | null | -1.63241 | -0.823486 | -0.21775 |
| YGR233C | PHO81 | -1.485611 | -0.980128 | -0.03703 |
| YDL042C | SIR2 | -1.420121 | -0.998536 | -0.09364 |
| YKR104W | null | -1.229861 | -0.567416 | 0.079281 |
| YOR066W | MSA1 | -1.373663 | -0.658872 | -0.06609 |
| YFL021W | GAT1 | -1.39381 | -0.739817 | 0.070213 |
| YOL020W | TAT2 | -1.611607 | -0.635395 | -0.02211 |
| YLR285C-A | null | -1.578821 | -0.490248 | 0.013808 |
| YBR296C | PHO89 | -1.466358 | -0.626254 | 0.239036 |
| YGR143W | SKN1 | -1.472033 | -0.868274 | 0.301284 |
| YBR196C-A | null | -1.686044 | -1.024496 | 0.365435 |
| YGR068C | ART5 | -1.308685 | -1.367768 | -0.25619 |
| YPR009W | SUT2 | -1.199259 | -1.578666 | -0.1354 |
| YPL066W | RGL1 | -1.00754 | -1.297962 | -0.02652 |
| YLR278C | null | -1.039162 | -1.355646 | 0.097341 |
| YBR291C | CTP1 | -1.350511 | -1.317544 | 0.568544 |
| YBL042C | FUI1 | -1.223819 | -0.91923 | 0.308592 |
| YOR342C | null | -1.000736 | -1.138968 | 0.459689 |
| YPR106W | ISR1 | -1.79978 | -0.483187 | -0.2268 |
| YOR084W | LPX1 | -1.799683 | -0.689908 | -0.43065 |
| YOR137C | SIA1 | -1.75522 | -0.752846 | -0.40568 |
| YIR019C | FLO11 | -1.986636 | -0.540304 | -0.32771 |
| YMR266W | RSN1 | -2.049499 | -0.570817 | -0.53056 |
| YDR072C | IPT1 | -1.593271 | 0.019658 | -0.28354 |
| YNR067C | DSE4 | -1.660394 | -0.136176 | -0.36397 |
| YGL028C | SCW11 | -1.830712 | -0.060328 | -0.32761 |
| YLR139C | SLS1 | -2.085937 | -0.41219 | 0.129681 |
| YOR383C | FIT3 | -1.95521 | -0.253077 | 0.131117 |
| YMR011W | HXT2 | -1.933943 | -0.067735 | 0.024425 |
| YGR249W | MGA1 | -1.655524 | -0.177218 | 0.18255 |
| YDR038C | ENA5 | -1.198654 | -0.029782 | -0.27106 |
| YMR155W | null | -1.220131 | -0.173316 | -0.18325 |
| YNL327W | EGT2 | -1.412944 | -0.034828 | -0.17192 |
| YDL195W | SEC31 | -1.348284 | -0.172596 | -0.38395 |
| YOL105C | WSC3 | -1.292077 | -0.338939 | -0.38492 |
| YAL021C | CCR4 | -1.257148 | -0.375177 | -0.42389 |
| YGL063W | PUS2 | -1.154045 | -0.379087 | -0.23419 |
| YOR081C | TGL5 | -1.149228 | -0.377802 | -0.29463 |
| YHR151C | MTC6 | -1.186217 | -0.269886 | -0.29443 |
| YCR048W | ARE1 | -1.105687 | -0.121713 | -0.47338 |
| YOR098C | NUP1 | -1.068292 | -0.063428 | -0.44489 |
| YOR149C | SMP3 | -1.034922 | -0.057169 | -0.43783 |
| YER154W | OXA1 | -1.039944 | -0.252443 | -0.41311 |
| YHR143W | DSE2 | -1.102243 | -0.245431 | -0.38771 |
| YEL017W | GTT3 | -1.29664 | -0.207355 | -0.66235 |
| YOL158C | ENB1 | -1.43844 | -0.320369 | -0.77348 |
| YOR068C | VAM10 | -1.052164 | -0.447297 | -0.52555 |
| YNL251C | NRD1 | -1.223235 | -0.360766 | -0.62714 |
| YBL004W | UTP20 | -1.193647 | -0.431908 | -0.60151 |
| YJL062W | LAS21 | -1.224755 | -0.486421 | -0.66804 |
| YOR160W | MTR10 | -1.199381 | -0.529148 | -0.52078 |
| YCR084C | TUP1 | -1.191201 | -0.64103 | -0.54665 |
| YHR042W | NCP1 | -1.132657 | -0.631171 | -0.55374 |
| YNL275W | BOR1 | -1.148786 | -0.62428 | -0.55425 |
| YJL198W | PHO90 | -1.249746 | -0.669753 | -0.43839 |
| YBL081W | null | -1.382356 | -0.512508 | -0.48926 |
| YPL056C | LCL1 | -1.305682 | -0.50234 | -0.49005 |
| YLR397C | AFG2 | -1.246947 | -0.564002 | -0.18438 |
| YPL041C | null | -1.197879 | -0.597596 | -0.27577 |
| YBR239C | ERT1 | -1.151174 | -0.694096 | -0.13321 |
| YMR310C | null | -1.088656 | -0.7018 | -0.10469 |
| YNL278W | CAF120 | -1.041454 | -0.441577 | -0.07463 |
| YKR053C | YSR3 | -1.066398 | -0.578892 | -0.05501 |
| YAL037W | null | -1.007443 | -0.678412 | 0.04718 |
| YAR029W | null | -1.053898 | -0.631209 | 0.287797 |
| YGL162W | SUT1 | -1.301283 | -0.314407 | -0.06362 |
| YOR359W | VTs1 | -1.213828 | -0.250429 | -0.0447 |
| YOR161C | PNS1 | -1.238826 | -0.276779 | -0.09367 |
| YPL076W | GPI2 | -1.312037 | -0.184898 | 0.136374 |
| YCL005W | LDB16 | -1.081465 | -0.277874 | 0.020287 |
| YJL216C | IMA5 | -1.05942 | -0.269345 | 0.081337 |
| YNL034W | null | -1.018354 | -0.190546 | -0.12339 |
| YMR013C | SEC59 | -1.08777 | -0.242867 | -0.1055 |
| YNL288W | CAF40 | -1.095138 | -0.264427 | -0.15225 |
| YBL101W-B | null | -1.014151 | -0.995121 | -0.19002 |
| YBR301W | PAU24 | -1.260484 | -0.925733 | -0.24493 |
| YNL204C | SPS18 | -1.175955 | -0.853385 | -0.27689 |
| YJL193W | null | -1.368008 | -0.862954 | -0.3361 |
| YDR143C | SAN1 | -1.293169 | -0.8678 | -0.39477 |
| YPR054W | SMK1 | -1.248613 | -1.072935 | -0.34598 |
| YNL291C | MID1 | -1.077109 | -0.769886 | -0.70305 |
| YDR538W | PAD1 | -1.040836 | -0.710045 | -0.72063 |
| YLR459W | GAB1 | -1.0176 | -0.644804 | -0.73487 |
| YDR404C | RPB7 | -1.108425 | -0.793386 | -0.61565 |
| YHR163W | SOL3 | -1.082184 | -0.831005 | -0.57453 |
| YOL052C | SPE2 | -1.04761 | -0.873968 | -0.6079 |
| YPR121W | THI22 | -1.021577 | -0.693926 | -0.36949 |
| YHR086W-A | null | -1.059386 | -0.784566 | -0.40747 |
| YNR055C | HOL1 | -1.113354 | -0.745037 | -0.50927 |
| YOR321W | PMT3 | -1.080195 | -0.7333 | -0.57796 |
| YKL078W | DHR2 | -1.047444 | -0.702301 | -0.58379 |
| YPL162C | null | -1.091111 | -0.657734 | -0.54618 |
| YBR086C | IST2 | -1.022796 | -0.622193 | -0.57988 |
| YMR274C | RCE1 | -1.413554 | -0.919902 | -0.55519 |
| YCR065W | HCM1 | -1.285848 | -0.962969 | -0.67737 |
| YEL065W | SIT1 | -1.326927 | -0.783409 | -0.60061 |
| YHR086W | NAM8 | -1.318066 | -0.818393 | -0.61668 |
| YGR281W | YOR1 | -1.281143 | -0.81167 | -0.52579 |
| YDL179W | PCL9 | -1.268798 | -0.745332 | -0.50394 |
| YLR141W | RRN5 | -1.303257 | -0.781836 | -0.77922 |
| YGR038W | ORM1 | -1.246203 | -0.704735 | -0.74587 |
| YLL031C | GPI13 | -1.30253 | -0.643208 | -0.68088 |
| YBR175W | SWD3 | -1.260419 | -1.254828 | -0.86015 |
| YER110C | KAP123 | -1.067729 | -1.093829 | -0.8948 |
| YGR131W | FHN1 | -1.038851 | -1.171217 | -0.9338 |
| YBR074W | PFF1 | -1.063322 | -1.206995 | -0.77965 |
| YMR277W | FCP1 | -1.041506 | -0.97864 | -0.84932 |
| YPR052C | NHP6A | -1.022073 | -0.965162 | -0.68902 |
| YLR380W | CSR1 | -1.085019 | -0.869708 | -0.74981 |
| YNL238W | KEX2 | -1.107759 | -0.93974 | -0.74767 |
| YPR128C | ANT1 | -1.14267 | -0.956547 | -0.6306 |
| YPL018W | CTF19 | -1.096172 | -0.982328 | -0.61275 |
| YCL002C | null | -1.095239 | -1.116041 | -0.62248 |
| YOR129C | AFI1 | -1.366272 | -1.293772 | -0.5899 |
| YNL080C | EOS1 | -1.293655 | -1.163043 | -0.53001 |
| YOR378W | AMF1 | -1.008824 | -0.401119 | -1.10451 |
| YGR146C-A | null | -1.037179 | -0.253796 | -1.17931 |
| YBL112C | null | -1.087553 | -0.294808 | -0.86761 |
| YEL007W | MIT1 | -1.450384 | -0.175074 | -1.11328 |
| YBL032W | HEK2 | -1.428298 | -0.549564 | -1.20662 |
| YMR055C | BUB2 | -1.32715 | -0.832963 | -0.8408 |
| YDL093W | PMT5 | -1.285507 | -0.896378 | -0.84831 |
| YLR381W | CTF3 | -1.309061 | -0.870505 | -0.88433 |
| YNR060W | FRE4 | -1.404624 | -0.994241 | -0.99664 |
| YJR124C | null | -1.421199 | -0.870187 | -0.82313 |
| YPL128C | TBF1 | -1.46835 | -0.971687 | -0.79808 |
| YML059C | NTE1 | -1.59741 | -0.877355 | -0.76925 |
| YDL160C | DHH1 | -1.460546 | -0.712223 | -0.80449 |
| YDR044W | HEM13 | -1.482312 | -0.685862 | -0.96316 |
| YDR414C | ERD1 | -1.115155 | -0.567869 | -1.01918 |
| YOR307C | SLY41 | -1.176199 | -0.6677 | -0.94427 |
| YEL042W | GDA1 | -1.208786 | -0.640769 | -1.01143 |
| YNL283C | WSC2 | -1.176128 | -0.87668 | -0.95097 |
| YOR067C | ALG8 | -1.246612 | -0.771258 | -1.00481 |
| YHR115C | DMA1 | -1.122361 | -0.725427 | -0.90375 |
| YDL048C | STP4 | -1.213949 | 1.304484 | 1.386136 |
| YCL068C | null | -1.523301 | 1.657408 | 1.603099 |
| YFR022W | ROG3 | -1.101124 | 1.820295 | 2.165995 |
| YCR108C | null | -1.06561 | 1.33408 | 2.05838 |
| YPL165C | SET6 | -2.289622 | 1.097893 | 2.20319 |
| YMR182C | RGM1 | -1.899901 | 1.352962 | 1.640581 |
| YDR247W | VHS1 | -1.800171 | 1.391117 | 1.98415 |
| YPL014W | null | -1.910601 | 1.698177 | 2.742215 |
| YER028C | MIG3 | -1.415682 | 1.989151 | 0.358269 |
| YOL011W | PLB3 | -1.942625 | 0.632071 | 1.215531 |
| YER188C-A | null | -2.006381 | 0.225677 | 0.750596 |
| YER088C | DOT6 | -1.230253 | 0.578579 | 0.870996 |
| YLR120C | YPS1 | -1.208372 | 0.519998 | 0.838073 |
| YBR298C | MAL31 | -1.420194 | 0.582458 | 0.997994 |
| YKL220C | FRE2 | -1.163377 | 0.038322 | 0.585177 |
| YOL136C | PFK27 | -1.330024 | 0.122457 | 0.50754 |
| YBL111C | null | -1.481711 | 0.309395 | 0.461409 |
| YHL040C | ARN1 | -1.378605 | 0.368258 | 0.793778 |
| YEL063C | CAN1 | -1.266327 | 0.260261 | 0.646736 |
| YOR273C | TPO4 | -1.594325 | 0.143492 | 0.763655 |
| YDR534C | FIT1 | -1.637009 | 0.208779 | 0.328522 |
| YOR381W | FRE3 | -1.59899 | 0.290977 | 0.265652 |
| YCR089W | FIG2 | -1.756856 | 0.166196 | 0.177603 |
| YPL057C | SUR1 | -1.782771 | 0.123331 | 0.541527 |
| YAL040C | CLN3 | -1.861156 | 0.015472 | 0.404578 |
| YLR152C | null | -1.285577 | -0.35867 | 0.317725 |
| YAL060W | BDH1 | -1.328006 | -0.452297 | 0.269904 |
| YBR021W | FUR4 | -1.114172 | -0.330952 | 0.547602 |
| YKL029C | MAE1 | -1.253982 | -0.332381 | 0.555527 |
| YMR272C | SCS7 | -1.212712 | -0.223203 | 0.405308 |
| YOL059W | GPD2 | -1.007211 | -0.056624 | 0.831813 |
| YGL256W | ADH4 | -1.093417 | -0.305503 | 0.710373 |
| YLL066W-B | null | -1.952956 | -0.33443 | 0.509491 |
| YAR010C | null | -1.799371 | -0.406109 | 0.645434 |
| YHL016C | DUR3 | -1.741483 | -0.492125 | 0.35648 |
| YBR196C-B | null | -1.369806 | -0.58939 | 0.657573 |
| YCL025C | AGP1 | -1.682185 | -0.331049 | 0.823672 |
| YDR205W | MSC2 | -1.001279 | 0.140402 | -0.1044 |
| YIL006W | YIA6 | -1.039886 | 0.124122 | -0.01684 |
| YLR337C | VRP1 | -1.096899 | 0.188038 | -0.00914 |
| YNL321W | VNX1 | -1.180766 | 0.100233 | -0.12292 |
| YMR120C | ADE17 | -1.211862 | 0.029671 | -0.08318 |
| YPL190C | NAB3 | -1.246992 | 0.338697 | -0.18057 |
| YOR140W | SFL1 | -1.290206 | 0.326112 | -0.15347 |
| YMR069W | NAT4 | -1.221564 | 0.089779 | 0.273154 |
| YKR102W | FLO10 | -1.275704 | 0.194769 | 0.316539 |
| YDR232W | HEM1 | -1.332911 | 0.045818 | 0.348183 |
| YDR246W-A | null | -1.440417 | 0.158604 | 0.091489 |
| YDL140C | RPO21 | -1.31474 | 0.154825 | 0.066087 |
| YBR020W | GAL1 | -1.311369 | 0.086624 | 0.107964 |
| YOR349W | CIN1 | -1.048967 | -0.045259 | 0.035903 |
| YAL020C | ATS1 | -1.068134 | -0.002847 | 0.125437 |
| YGR079W | null | -1.15463 | -0.053631 | 0.260586 |
| YER152C | null | -1.022108 | -0.125026 | 0.246497 |
| YAL063C | FLO9 | -1.067861 | -0.071818 | 0.206601 |
| YKL187C | FAT3 | -1.036015 | 0.178003 | 0.196359 |
| YLR116W | MSL5 | -1.06763 | 0.179945 | 0.243781 |
| YKL198C | PTK1 | -1.096863 | 0.259249 | 0.20616 |
| YDR160W | SSY1 | -1.019812 | 0.148067 | 0.308233 |
| YER060W | FCY21 | -1.029224 | 0.203936 | 0.351275 |
| YGL178W | MPT5 | -1.179571 | 0.479767 | 0.359187 |
| YDR438W | THI74 | -1.258853 | 0.579677 | 0.370009 |
| YLR099C | ICT1 | -1.13923 | 0.600478 | 0.329931 |
| YLR403W | SFP1 | -1.212804 | 0.429344 | 0.237323 |
| YJL094C | KHA1 | -1.017959 | 0.514731 | 0.080093 |
| YMR008C | PLB1 | -1.103382 | 0.676622 | 0.553909 |
| YGR289C | MAL11 | -1.084481 | 0.696059 | 0.692787 |
| YPL015C | HST2 | -1.078514 | 1.148372 | 0.591381 |
| YHR094C | HXT1 | -1.578098 | 0.964523 | 0.494114 |
| YGR023W | MTL1 | -1.551499 | 0.762923 | 0.690967 |
| YMR291W | TDA1 | -1.2614 | 0.269249 | 1.804324 |
| YGR032W | GSC2 | -1.242827 | 0.236364 | 1.776908 |
| YJL079C | PRY1 | -1.443107 | 0.230725 | 1.836415 |
| YBR067C | TIP1 | -1.191979 | 0.6789 | 2.078076 |
| YHL026C | null | -1.244398 | 0.708651 | 1.464342 |
| YGR121C | MEP1 | -1.02356 | 0.688905 | 1.588961 |
| YER053C | PIC2 | -1.617607 | 0.204335 | 1.427363 |
| YJL212C | OPT1 | -1.53264 | 0.103232 | 1.427167 |
| YER158C | null | -1.423766 | 0.170882 | 1.213202 |
| YPL036W | PMA2 | -1.151877 | -0.145902 | 1.544101 |
| YLR142W | PUT1 | -1.002157 | 0.184707 | 1.238826 |
| YOR071C | NRT1 | -1.17284 | -0.074057 | 1.26817 |
| YGR260W | TNA1 | -1.147552 | 0.084828 | 1.242491 |
| YFL014W | HSP12 | -1.292276 | 0.237658 | 2.488439 |
| YPL092W | SSU1 | -1.065039 | -0.681846 | 0.960938 |
| YBL005W-A | null | -1.17081 | -0.719291 | 0.961157 |
| YNL024C | null | -1.26106 | -0.90903 | 1.338866 |
| YNR014W | null | -1.922514 | -0.605679 | 1.777716 |
| YPR160W | GPH1 | -1.099576 | -0.583531 | 2.009052 |
| YOL104C | NDJ1 | -1.647413 | -1.534572 | 1.433746 |
| YDR222W | null | -2.011217 | -1.458479 | 0.856427 |
| YCR018C | SRD1 | -1.838414 | -2.480437 | 1.389827 |
| YHR033W | null | -1.916852 | -1.503132 | 2.110011 |
| YOL052C-A | DDR2 | -1.365443 | -0.876442 | 3.35148 |
| YMR081C | ISF1 | -1.715576 | -1.687342 | 3.543186 |
| YHR092C | HXT4 | -3.350206 | 0.671244 | 0.82595 |
| YDL039C | PRM7 | -2.859618 | 0.584578 | 0.03101 |
| YDL038C | null | -3.677256 | 0.345895 | -0.19471 |
| YBR238C | null | -2.643991 | -0.768649 | -0.05108 |
| YLR466C-B | null | -2.556635 | -0.702605 | 0.286957 |
| YOR394C-A | null | -2.463799 | -0.091134 | 0.527107 |
| YLR466C-B | null | -2.4757 | -0.355529 | 0.395135 |
| YLL066W-B | null | -2.254415 | -0.214519 | 0.728592 |
| YJL078C | PRY3 | -2.463906 | 0.045136 | 0.17965 |
| YDR342C | HXT7 | -2.174849 | -0.87236 | 1.142651 |
| YER067W | RGI1 | -2.320877 | -1.049696 | 1.041793 |
| YOR092W | ECM3 | -2.179419 | -0.617264 | 0.945413 |
| YLR121C | YPS3 | -2.161079 | -0.657103 | 0.534248 |
| YCR097W | HMRA1 | -5.067512 | -0.8165 | 0.728803 |

**Table S3:**
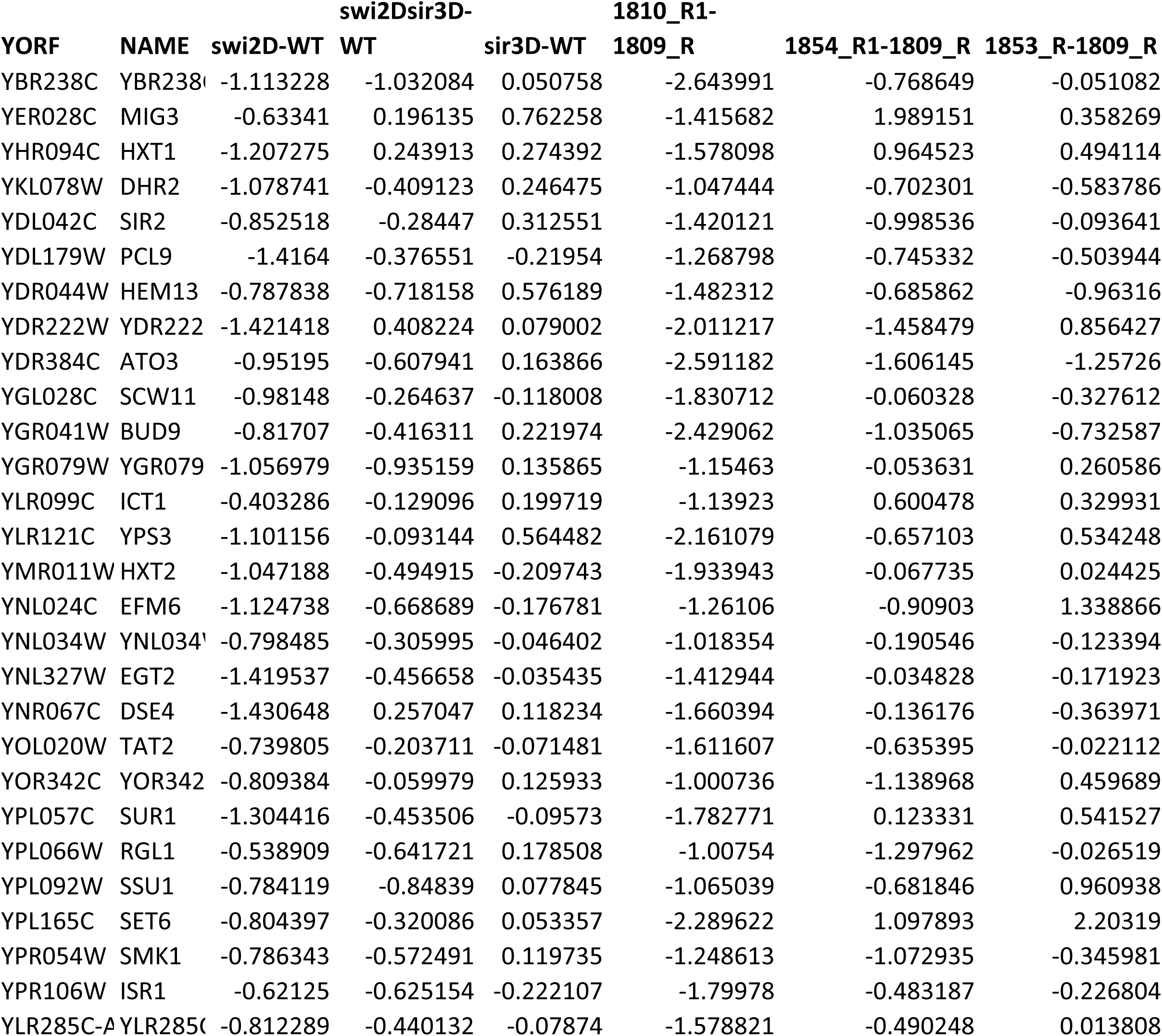
List of Group 1 genes common in the KO and AA datasets. swi2D = swi2 null mutant; swi2Dsir3D = swi2 and sir3 null mutant, sir3D = sir3 null mutant, WT = Wildtype, 1810 = SWI2 Anchor away; 1854= SWI2 Anchor away sir3 null mutant; 1853 = sir3 null in Anchor away background; 1809 = Wildtype Anchor away background. _R = with Rapamycin

**Table S4:**
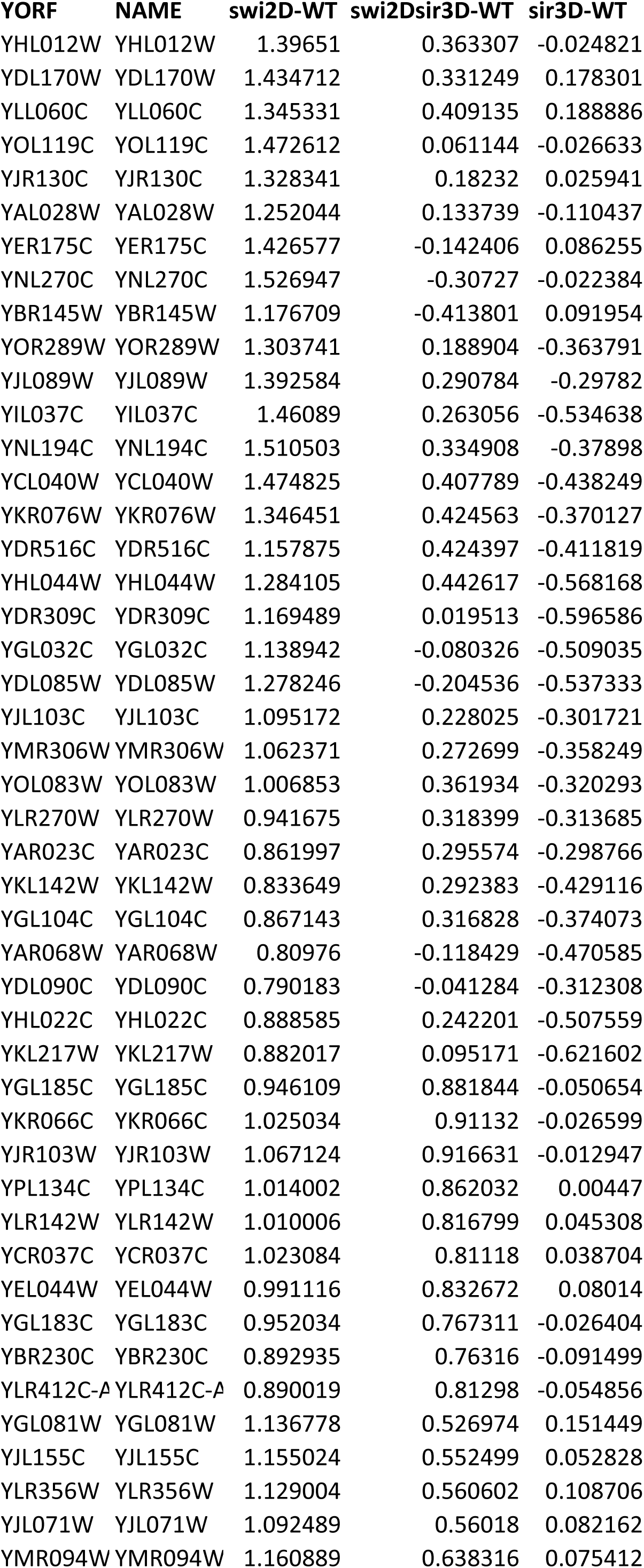
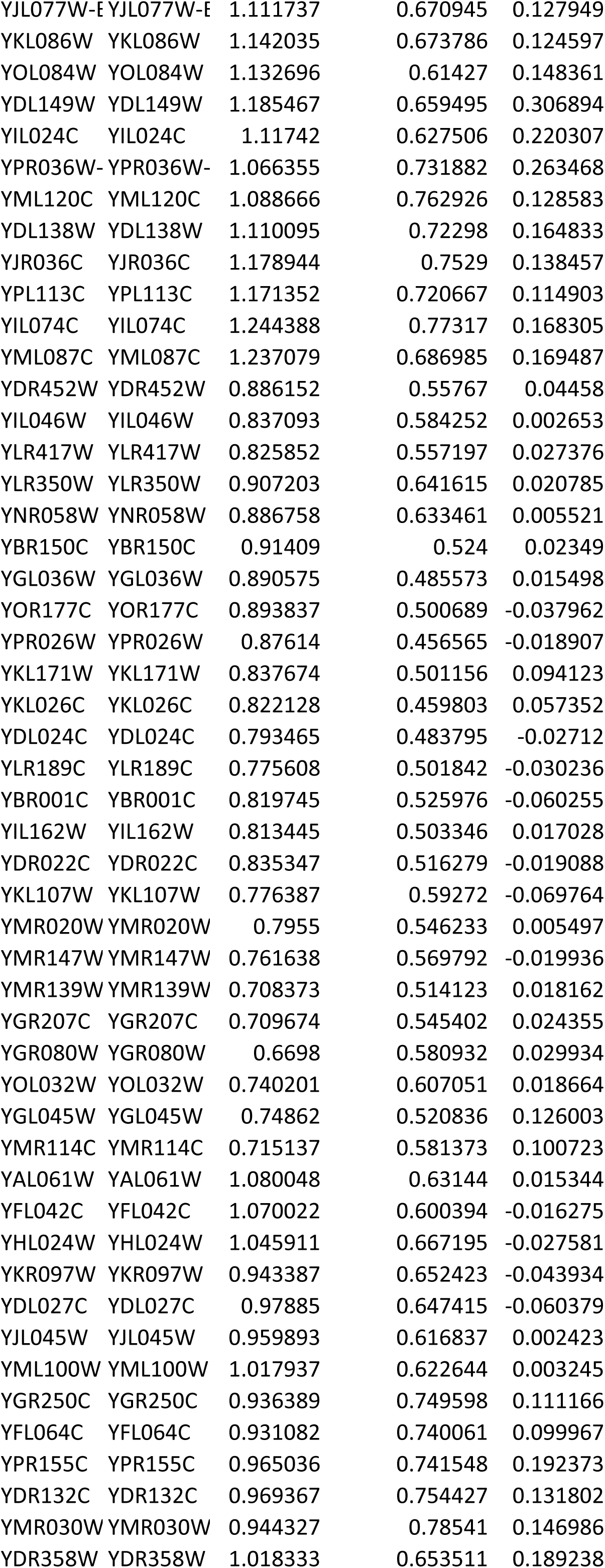
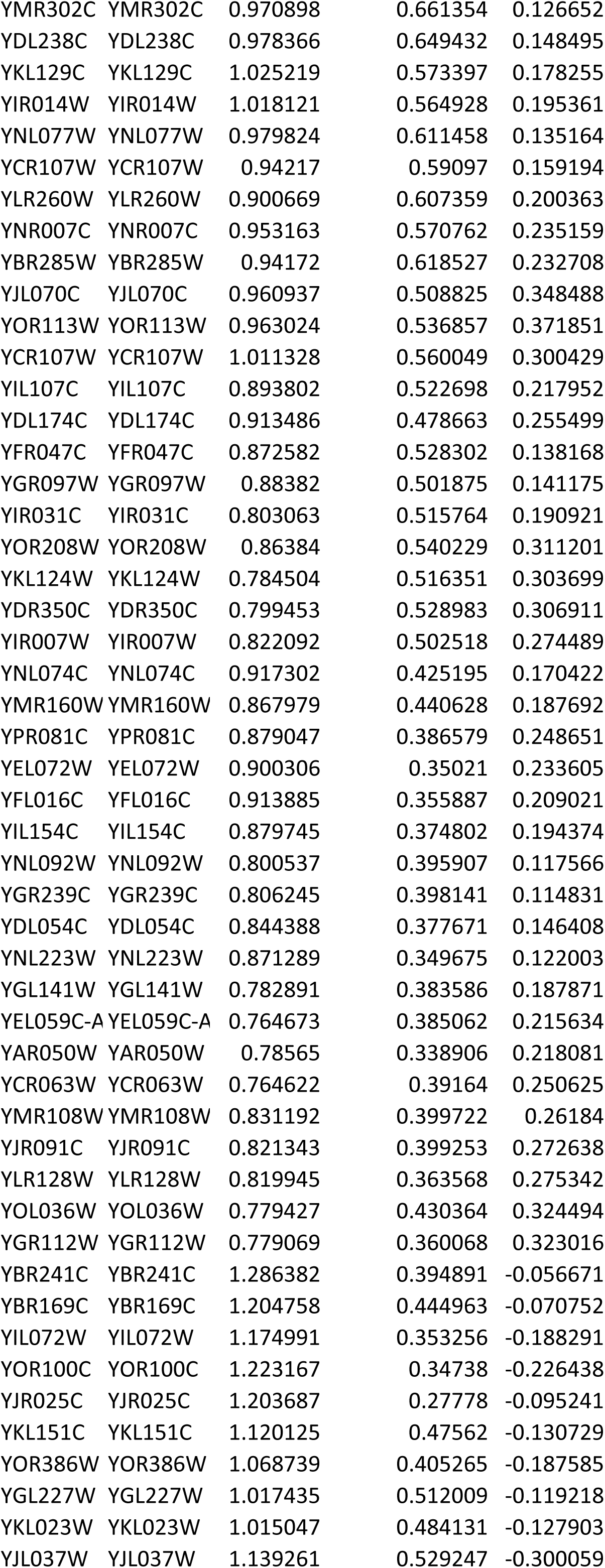
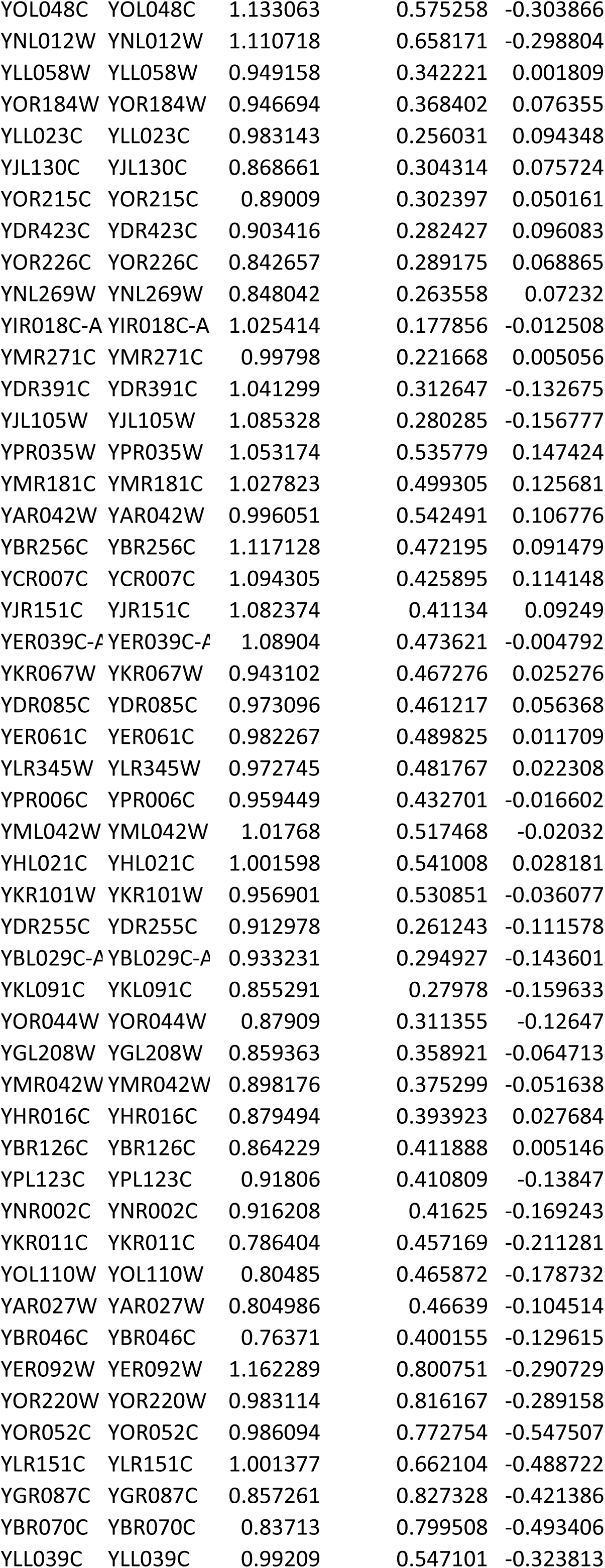
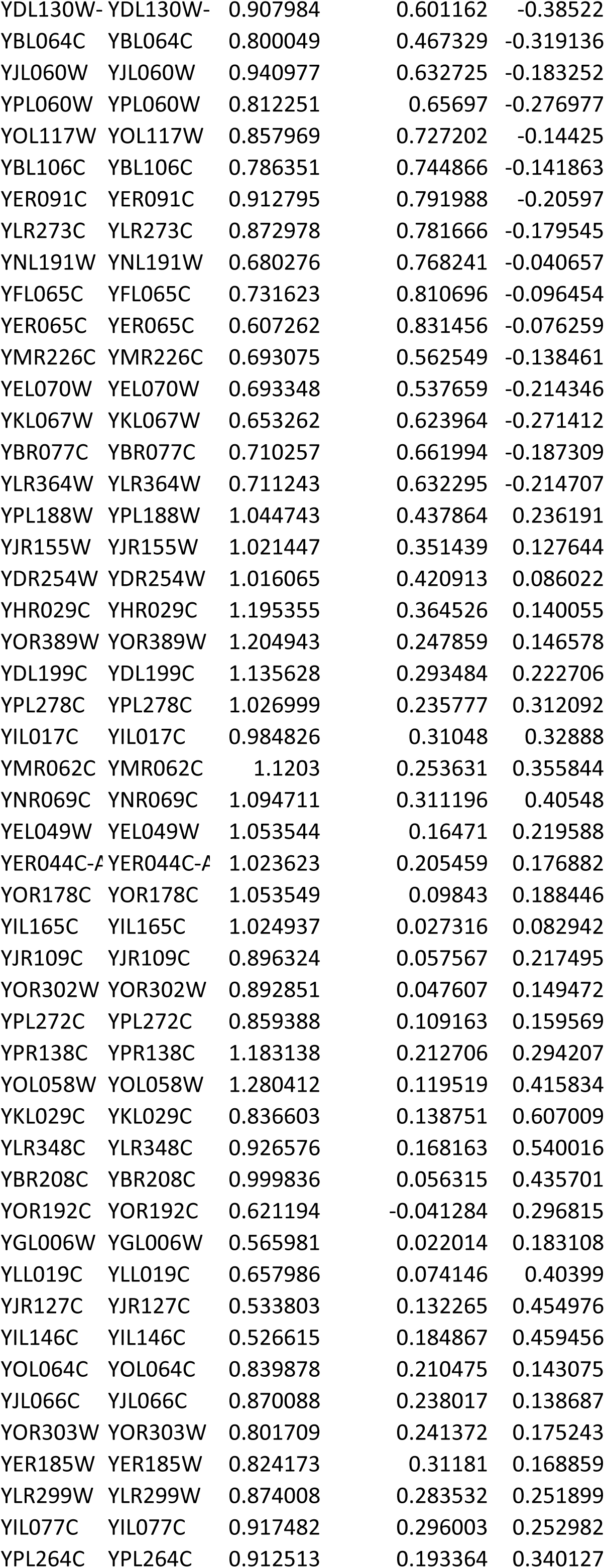
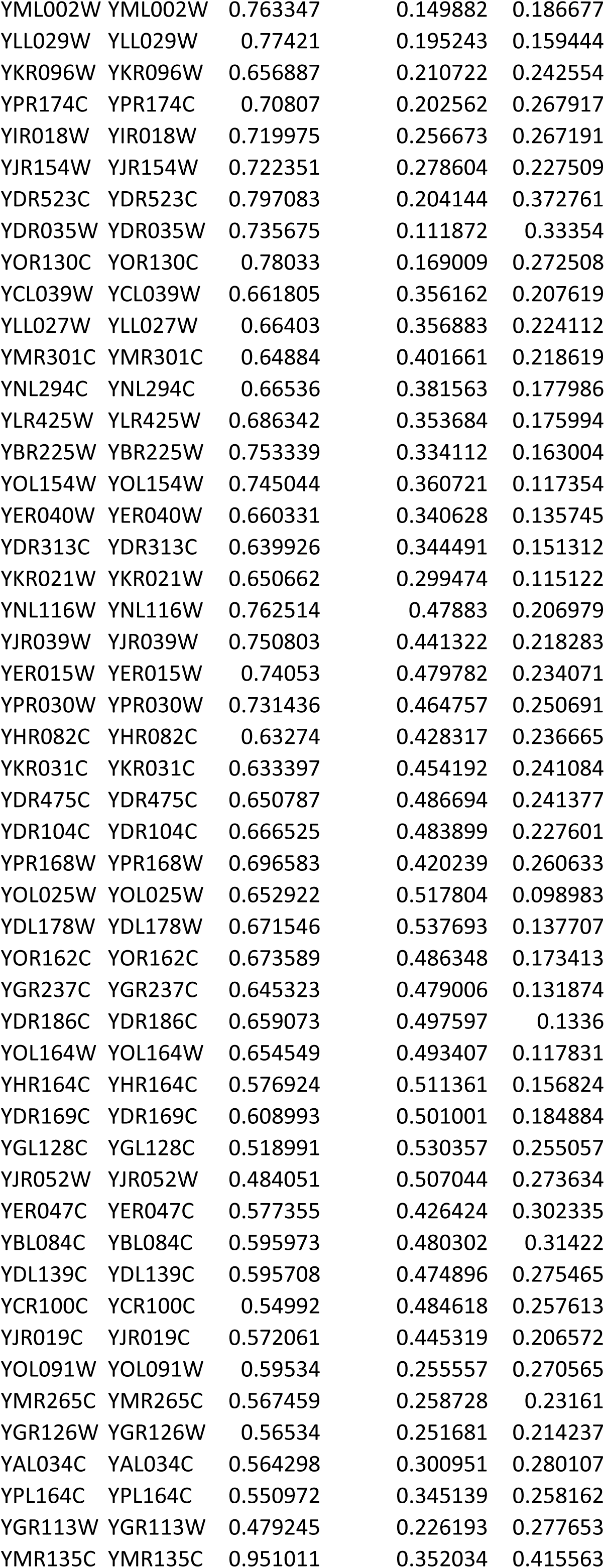
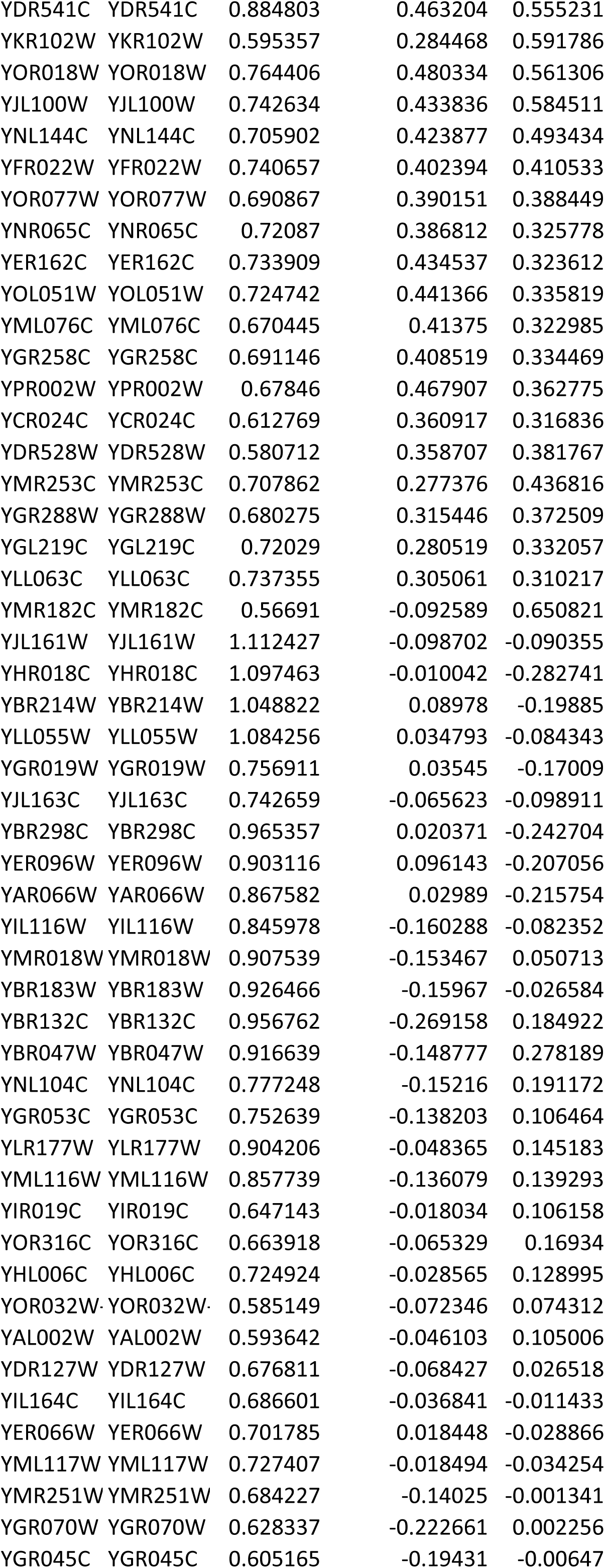
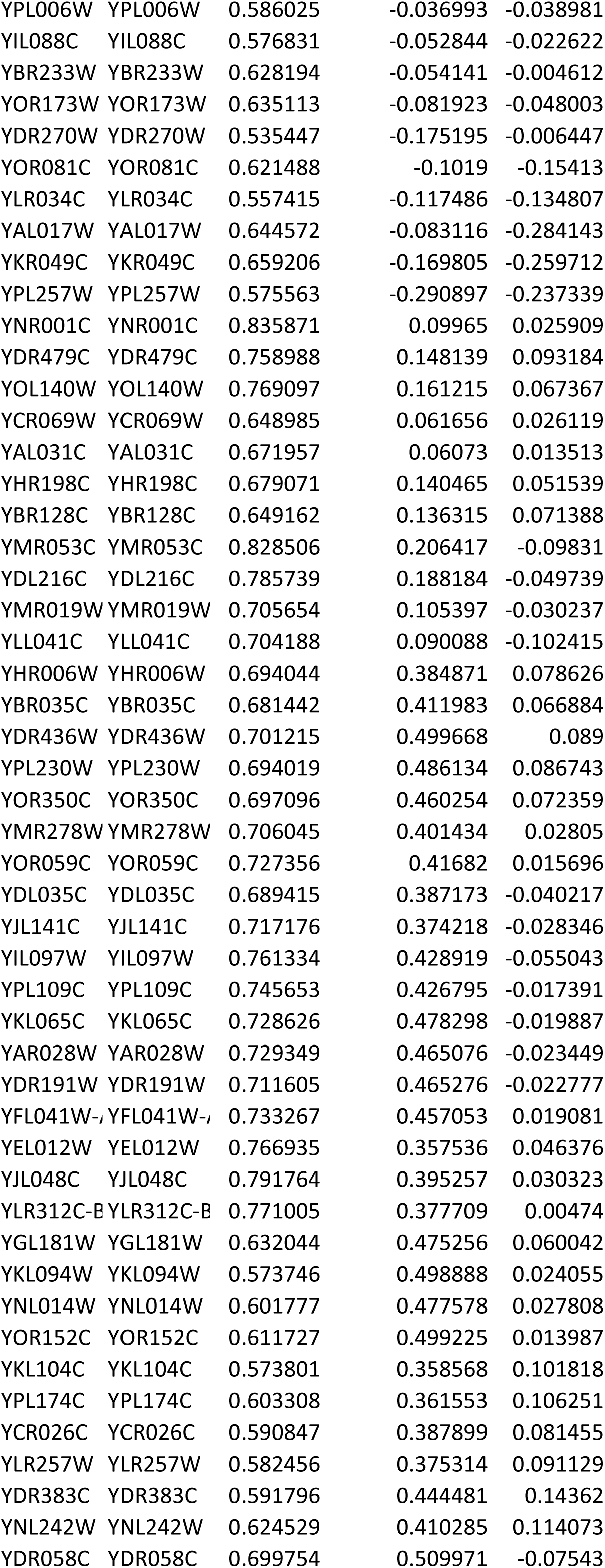
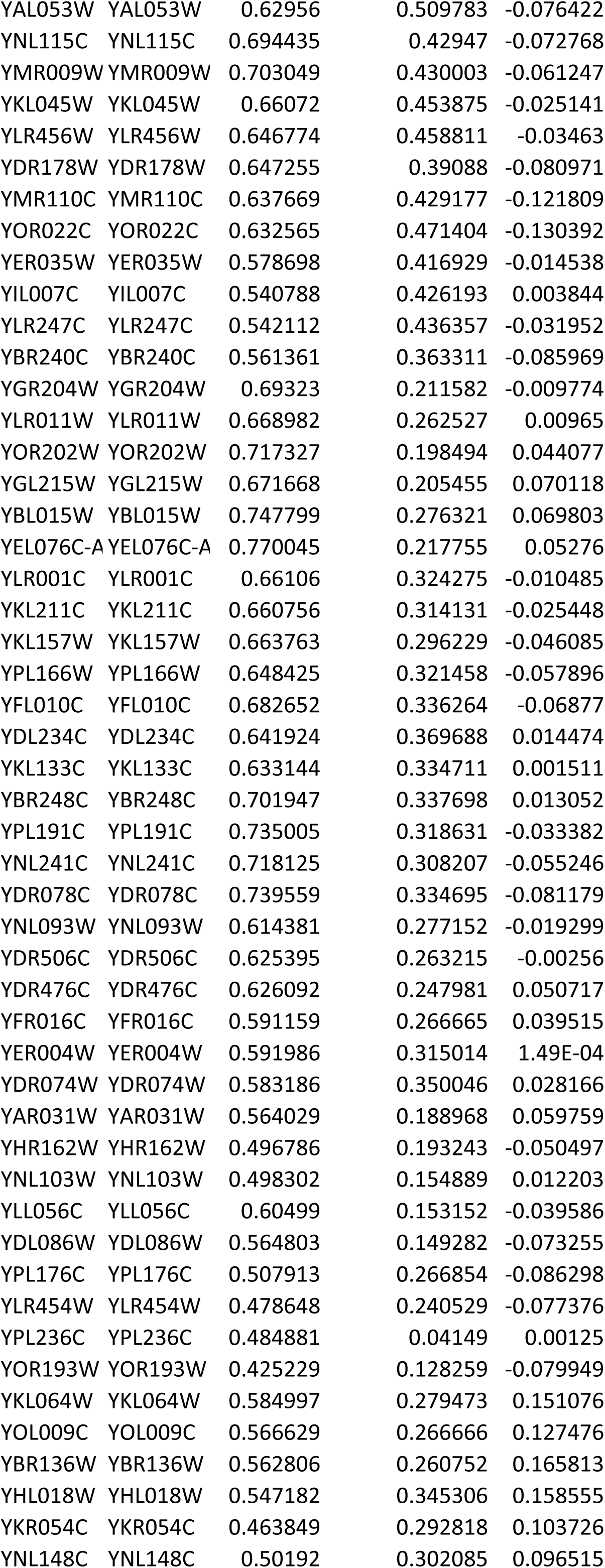
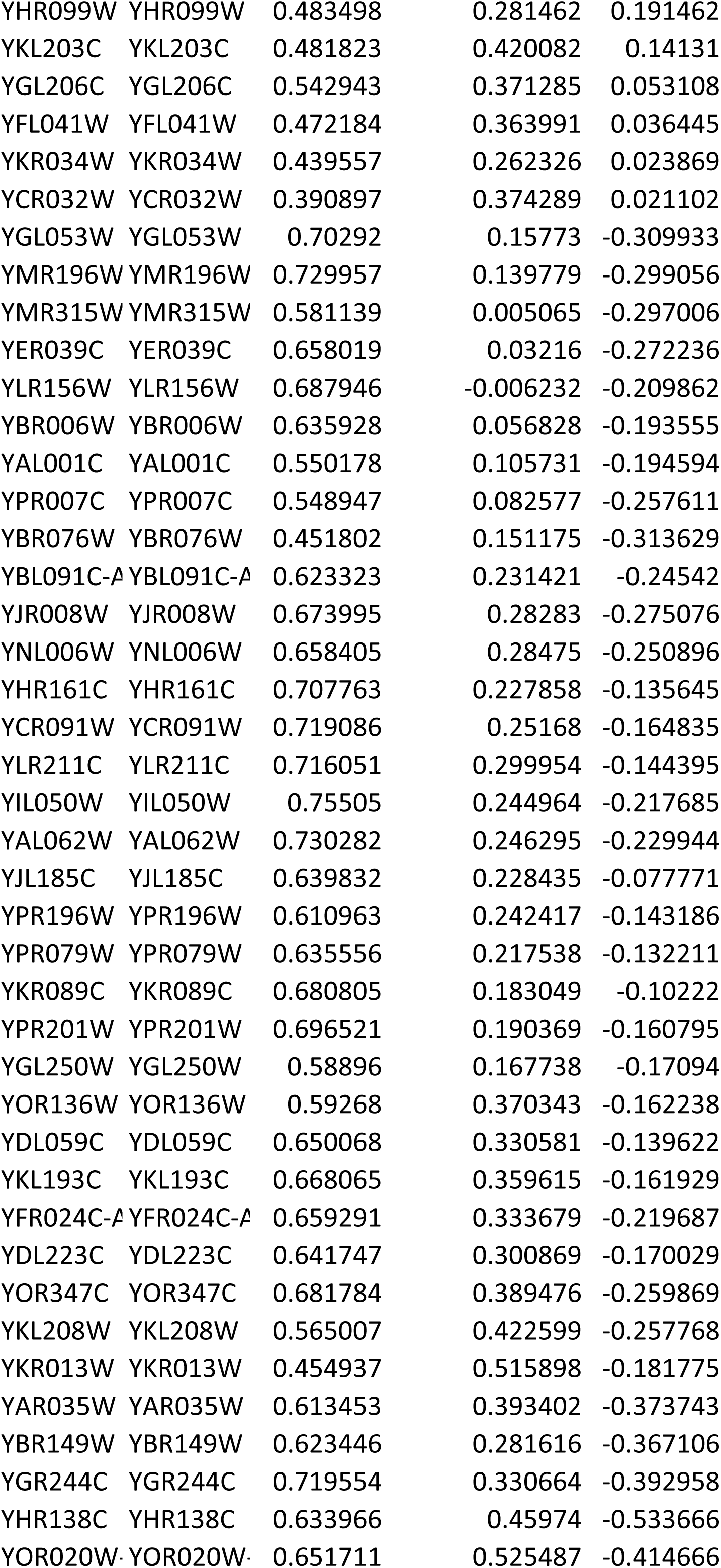
List of Group 2_KO genes. swi2D = swi2 null mutant; sir3D = sir3 null mutant, WT = Wildtype

**Table S5:**
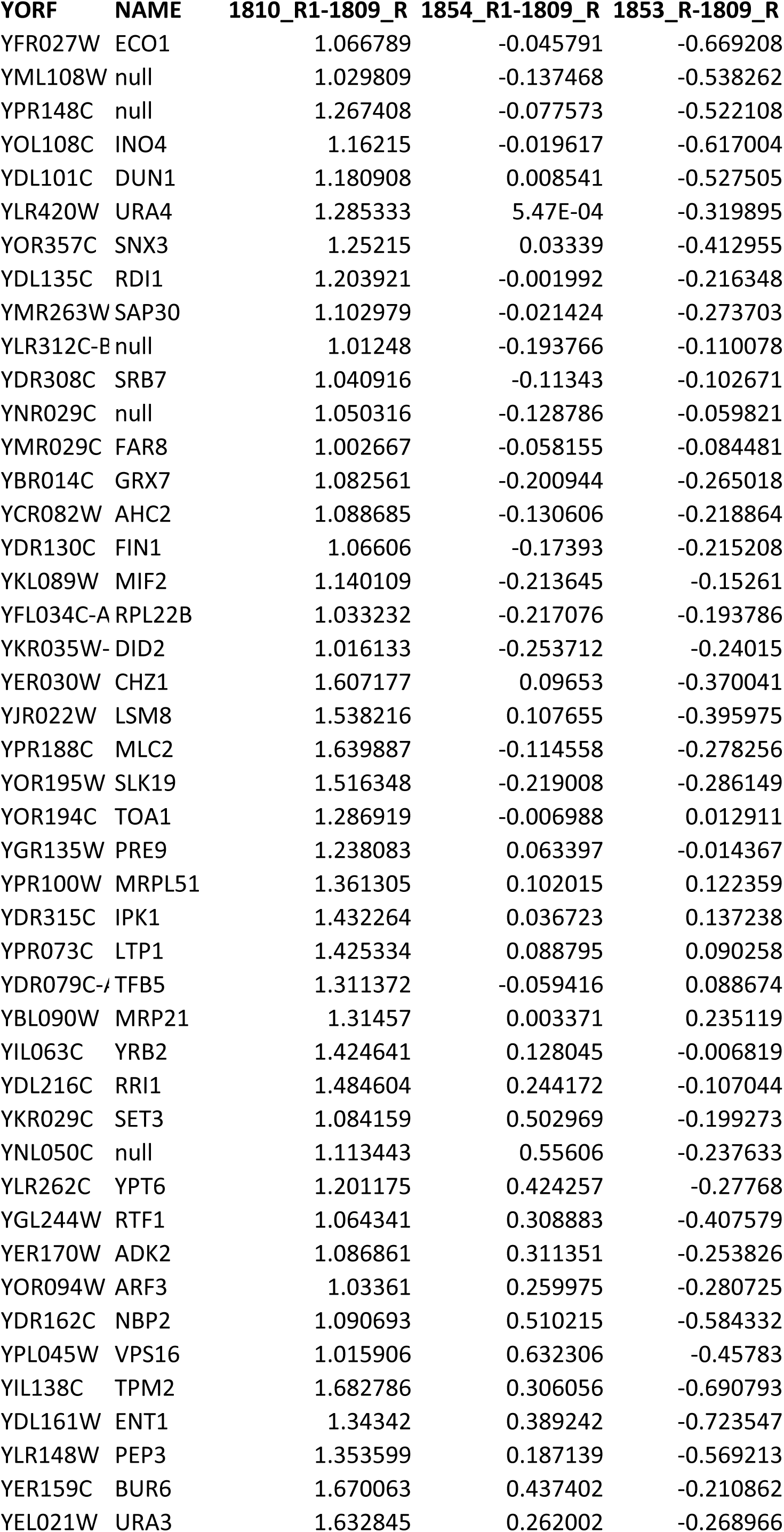
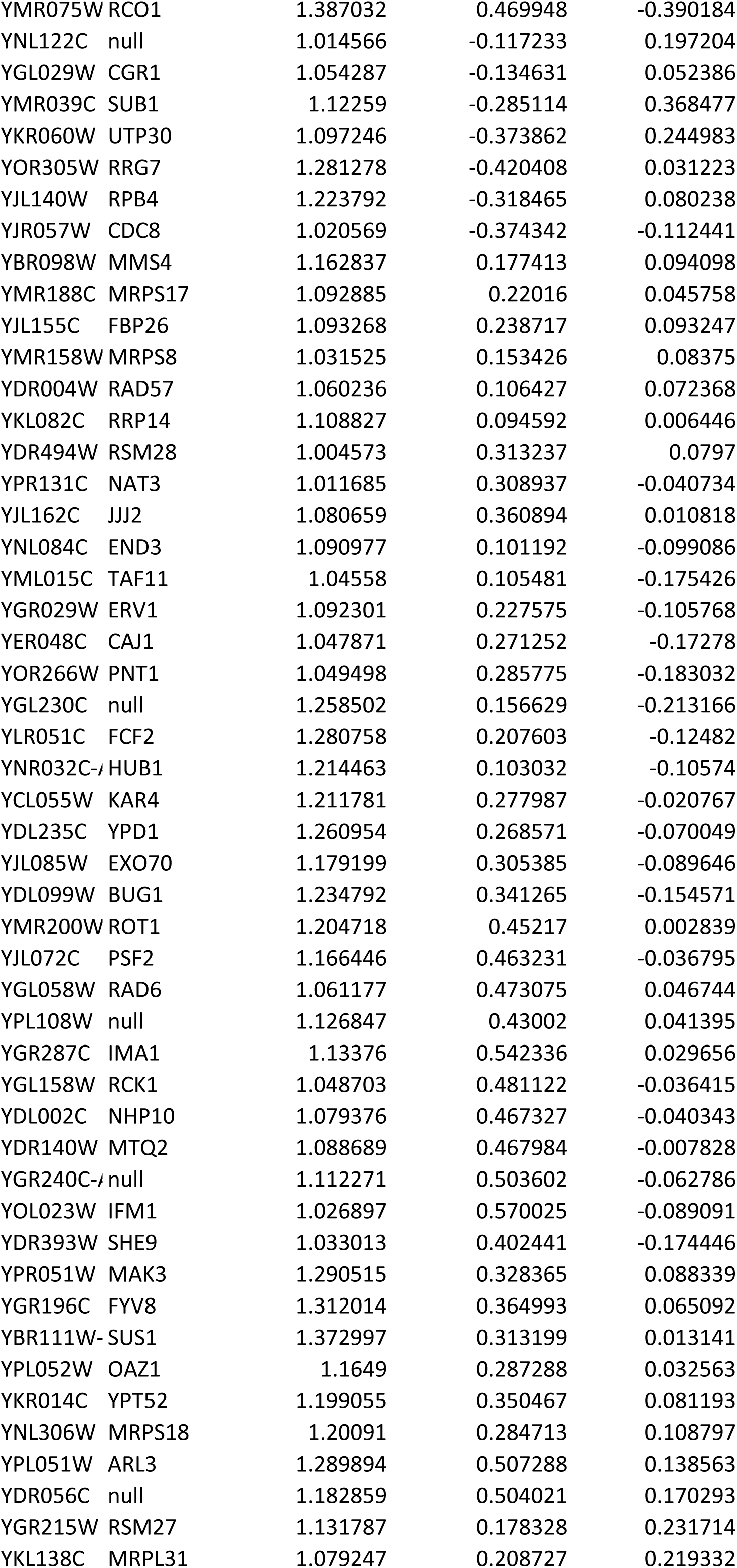
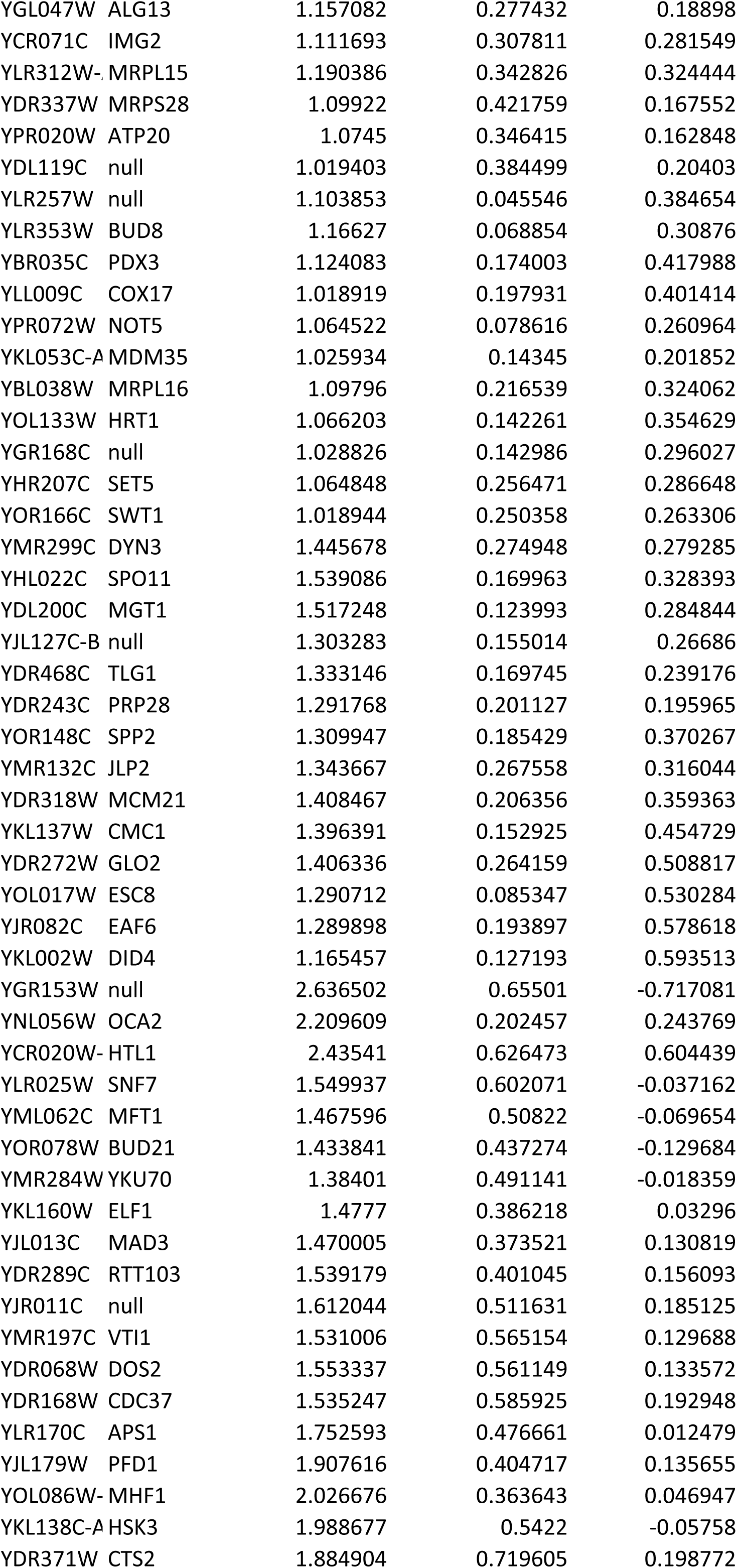
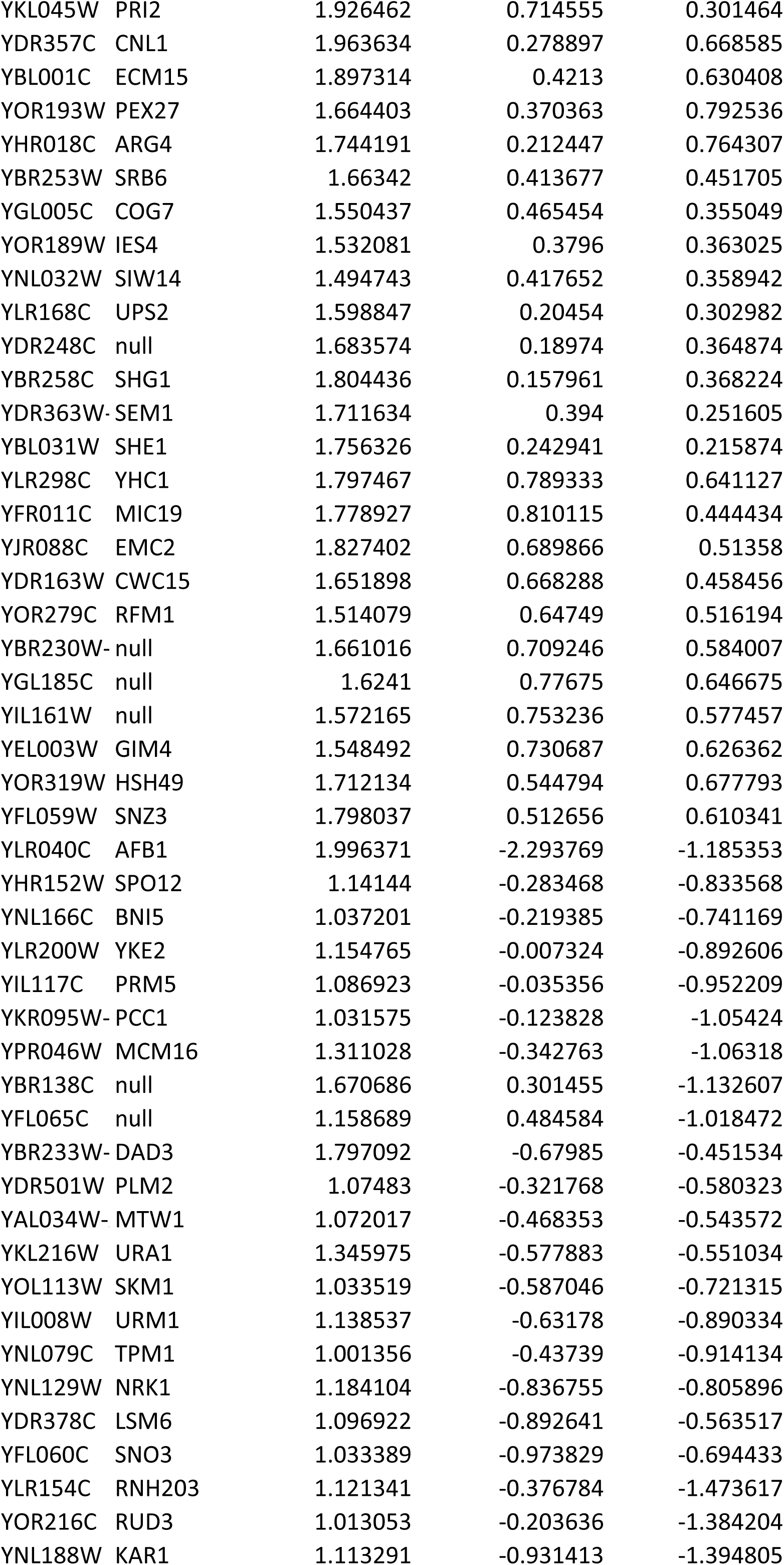
List of Group 2_AA genes. 1810 = SWI2 Anchor away; 1854 = SWI2 Anchor away sir3 null mutant; 1853 = sir3 null in Anchor away background; 1809 = Wildtype Anchor away background. _R = with Rapamycin

**Table S6: Complete table of RMA values from the KO datasets for all genes**

**Table S7: Complete table of RMA values from the AA datasets for all genes**

